# Comprehensive comparative assessment of the *Arabidopsis thaliana* MLO2-calmodulin interaction by various *in vitro* and *in vivo* protein-protein interaction assays

**DOI:** 10.1101/2023.01.25.525488

**Authors:** Kira von Bongartz, Björn Sabelleck, Anežka Baquero Forero, Hannah Kuhn, Franz Leissing, Ralph Panstruga

## Abstract

Mildew resistance locus o (MLO) proteins are heptahelical integral membrane proteins of which some isoforms act as susceptibility factors for the fungal powdery mildew pathogen. In many angiosperm plant species, loss-of-function *mlo* mutants confer durable broad-spectrum resistance against the powdery mildew disease. Barley Mlo is known to interact *via* a cytosolic carboxyl-terminal domain with the intracellular calcium sensor calmodulin (CAM) in a calcium-dependent manner. Site-directed mutagenesis has revealed key amino acid residues in the barley Mlo calcium-binding domain (CAMBD) that, when mutated, affect the MLO-CAM association. We here tested the respective interaction between *Arabidopsis thaliana* MLO2 and CAM2 using seven different types of *in vitro* and *in vivo* protein-protein interaction assays. In each assay, we deployed a wild-type version of either the MLO2 carboxyl terminus (MLO2^CT^), harboring the CAMBD, or the MLO2 full-length protein and corresponding mutant variants in which two key residues within the CAMBD were substituted by non-functional amino acids. We focused in particular on the substitution of two hydrophobic amino acids (LW/RR mutant) and found in most protein-protein interaction experiments reduced binding of CAM2 to the corresponding MLO2/MLO2^CT^ LW/RR mutant variants in comparison to the respective wild-type versions. However, the Ura3-based yeast split-ubiquitin system and *in planta* bimolecular fluorescence complementation (BiFC) assays failed to indicate reduced CAM2 binding to the mutated CAMBD. Our data shed further light on the interaction of MLO and CAM proteins and provide a comprehensive comparative assessment of different types of protein-protein interaction assays with wild-type and mutant versions of an integral membrane protein.

## Introduction

Interactions between biomolecules are key for all processes of life. Of particular interest are intermolecular contacts between proteins as these macromolecules are multifunctional cellular workhorses. Proteins get in contact with each other *via* surfaces formed by their respective amino acid residue side chains. Mutual attachment between them relies on combinations of reversible ionic interactions and hydrogen bonds, as well as van der Waals forces and other types of hydrophobic bondings that form between the amino acids of the interacting proteins (Erijman *et al*. 2014). Depending on the identity and number of amino acid residues involved, protein-protein interactions can be stable or transient, strong or weak (Erijman *et al*. 2014). They can be modulated by additional factors such as the composition of the solvent medium (Ebel 2007), the occurrence of post-translational protein modifications (Duan and Walther 2015) and/or the participation of additional (competing or supporting) binding partners. Due to their importance in biological processes, a plethora of methods has been developed to study protein-protein interactions *in vitro* and *in vivo*. Not surprisingly, each method has its specific advantages and disadvantages (Xing *et al*. 2016). Accordingly, no consensus has been reached so far regarding a commonly accepted “gold standard” for probing protein-protein interactions.

Mildew resistance locus o (MLO) proteins are integral membrane proteins that in most cases have seven predicted membrane-spanning domains, an extracellular/luminal N-terminus, and a cytosolic C-terminus. Although distantly related members have been identified in algae and some oomycetes, the protein family expanded predominantly within the embryophytes (land plants; Kusch *et al*. 2016). In seed plants, for example, approximately 10-20 paralogs exist per species. The founding and eponymous member of the family is barley Mlo. The barley *Mlo* gene was initially discovered as a locus that in its wild-type allelic form confers susceptibility to the fungal powdery mildew disease. Conversely, recessively inherited loss-of-function *mlo* mutants provide exceptionally durable broad-spectrum resistance to the pathogen (Jørgensen 1992). This mutant phenotype is largely conserved between angiosperm plants that can be hosts for powdery mildew fungi (Kusch and Panstruga 2017). Accordingly, *mlo* mutants, especially in barley, are of great agricultural and economical importance (Lyngkjær *et al*. 2000). In some plant species, however, multiple *Mlo* co-orthologs exist. In the dicotyledonous reference plant *Arabidopsis thaliana*, for example, genes *MLO2*, *MLO6* and *MLO12* are the co-orthologs of barley *Mlo* and modulate powdery mildew susceptibility in a genetically unequal manner. Of these three genes, *MLO2* is the main player in the context of powdery mildew disease (Consonni *et al*. 2006).

Extensive genetic studies, mostly conducted in *A. thaliana*, revealed that other members of the MLO family contribute to different biological processes. For example, *MLO4* and *MLO11* are implicated in root thigmomorphogenesis (Bidzinski *et al*. 2014; Chen *et al*. 2009), *MLO7* governs pollen tube reception at the female gametophyte (Kessler *et al*. 2010; Jones *et al*. 2017), *MLO5*, *MLO9* and *MLO15* modulate pollen tube guidance in response to ovular signals (Meng *et al*. 2020), and MLO3, similar to MLO2 (Consonni *et al*. 2010; Consonni *et al*. 2006), controls the timely onset of leaf senescence (Kusch *et al*. 2019). Moreover, MLO2 acts also negative regulator of sensitivity to extracellular reactive oxygen species (Cui *et al*. 2018).

Apart from its predicted, and in the case of barley Mlo experimentally validated, heptahelical membrane topology (Devoto *et al*. 1999), MLO proteins share a framework of conserved amino acid residues. These include four luminally/extracellularly positioned cysteine residues that are predicted to form two disulfide bridges (Elliott *et al*. 2005), and some short peptide motifs (Devoto *et al*. 1999; Kusch *et al*. 2016; Panstruga 2005) dispersed throughout the protein. A further common feature is the existence of a predicted and in part experimentally validated binding domain for the small (∼18 kDa molecular mass) cytosolic calcium sensor protein, calmodulin (CAM). This stretch is comprised of approximately 15-20 amino acids and is located at the proximal end of the C-terminal cytoplasmic tail region of Mlo proteins (Kim *et al*. 2002a; Kim *et al*. 2002b). It is supposed to form an amphiphilic a-helix, with (positively charged) hydrophilic residues primarily located on one side of the helix and (uncharged) hydrophobic residues on the other, thereby forming a CAM-binding domain (CAMBD). Calcium-induced conformational changes in the four EF hands of CAM allow for the binding of the calcium sensor protein to the MLO CAMBD. This was experimentally evidenced by yeast-based interaction assays (Kim *et al*. 2002b; Zhu *et al*. 2021; Yu *et al*. 2019), *in vitro* binding studies (Kim *et al*. 2002a; Kim *et al*. 2002b), co-immunoprecipitation experiments (Kim *et al*. 2014), as well as *in planta* Luciferase Complementation Imaging (LCI) (Zhu *et al*. 2021; Yu *et al*. 2019), Bimolecular fluorescence complementation (BiFC) (Zhu *et al*. 2021; Kim *et al*. 2014; Yu *et al*. 2019) and Fluorescence Resonance Energy Transfer (FRET) assays (Bhat *et al*. 2005), using combinations of different Mlo and CAM/CAM-like proteins (CMLs) from various plant species.

Site-directed mutagenesis has revealed the importance of key hydrophobic amino acid residues within the CAMBD for the establishment of the MLO-CAM interaction. Amino acid substitutions of these essential residues with positively charged arginines largely prevented the calcium-dependent binding of CAM to the CAMBDs of barley and rice MLO proteins (Kim *et al*. 2002a; Kim *et al*. 2002b). The reduction in CAM binding has consequences for the physiological role of barley Mlo: Respective mutations in the CAMBD lower the susceptibility-conferring capacity of the protein, as revealed by single cell expression experiments (Kim *et al*. 2002b). Whether similar site-directed mutations would also affect the CAMBD of *A. thaliana* MLO2, which like barley Mlo is implicated in the modulation of powdery mildew susceptibility (Consonni *et al*. 2006), remained elusive.

We here explored the interaction between *A. thaliana* MLO2 and the CAM isoform CAM2 using seven different assays to visualize protein-protein interactions. These comprise both *in vitro* and *in vivo* approaches, are based on either the isolated MLO2 carboxyl terminus (MLO2^CT^) or the full-length MLO2 protein, and rely on entirely different types of signal output. We found that except for the classical yeast two-hybrid (Y2H) approach, each method indicated interaction between MLO2/MLO2^CT^ and CAM2. We further created several single amino acid substitution mutant variants within the MLO2 CAMBD and tested these for interaction with CAM2. We focused in particular on the substitution of two key hydrophobic amino acids by arginines (LW/RR mutant). We found that most of the protein assays that indicate interaction between MLO2/MLO2^CT^ and CAM2 also faithfully specified reduced binding of CAM2 to the respective LW/RR mutant variants. Our data offer a detailed characterization of the MLO2 CAMBD and provide a showcase for the comparative assessment of different *in vitro* and *in vivo* protein-protein interaction assays with wild-type (WT) and mutant versions of an integral membrane protein.

## Results

### *In silico* analysis of the predicted MLO2^CT^ and its associated CAMBD

Similar to other MLO proteins (Kusch *et al*. 2016; Devoto *et al*. 1999), the *in silico* determined membrane topology of MLO2 (Arabidopsis Genome Initiative identifier At1g11310) comprises seven transmembrane domains, an extracellular/luminal N-terminus, and a cytoplasmic C-terminus (MLO2^CT^; **Figure 1A**). We performed a prediction of the three-dimensional structure of the cytoplasmic MLO2^CT^ by AlphaFold (https://alphafold.ebi.ac.uk/; Jumper *et al*. 2021). This revealed the presence of an α-helical region between amino acids R^451^ and K^468^, spanning the presumed CAMBD, and otherwise the absence of extended structural folds, suggesting that the MLO2^CT^ is largely intrinsically disordered (**Figure 1B**). This outcome agrees well with the calculation by PONDR-FIT (http://original.disprot.org/pondr-fit.php; Xue *et al*. 2010), a meta-predictor of intrinsically disordered protein regions, which indicates a high disorder tendency for the MLO2^CT^ (approximately after residue 475; **Figure 1C**). The combined *in silico* analysis using AlphaFold and PONDR-FIT suggests that the proposed CAMBD is the main structured segment of the MLO2^CT^. We subjected the proposed α-helical region, covering the presumed CAMBD of MLO2, to helical wheel projection by pepwheel (https://www.bioinformatics.nl/cgi-bin/emboss/pepwheel). We found that, as expected for a genuine CAMBD, this stretch of the MLO2^CT^ is estimated to form an amphiphilic α-helix, with hydrophilic amino acids primarily located on one side of the helix and hydrophobic residues mainly occupying the opposite side (**Figure 1D**). A comparison of the helical wheel projections of the predicted MLO2 CAMBD with the CAMBD of barley Mlo revealed several conserved amino acid positions among the two proteins (**Supplemental File 1** and **Supplemental Figure 1**). These included, amongst others, invariant leucine and tryptophan residues (L^18^ and W^21^ in MLO2^CT^; corresponding to L^456^ and W^459^ in full-length MLO2) that were previously shown to be essential for CAM binding to the CAMBD in barley and rice MLO proteins (Kim *et al*. 2002a; Kim *et al*. 2002b).

**Figure 1.**
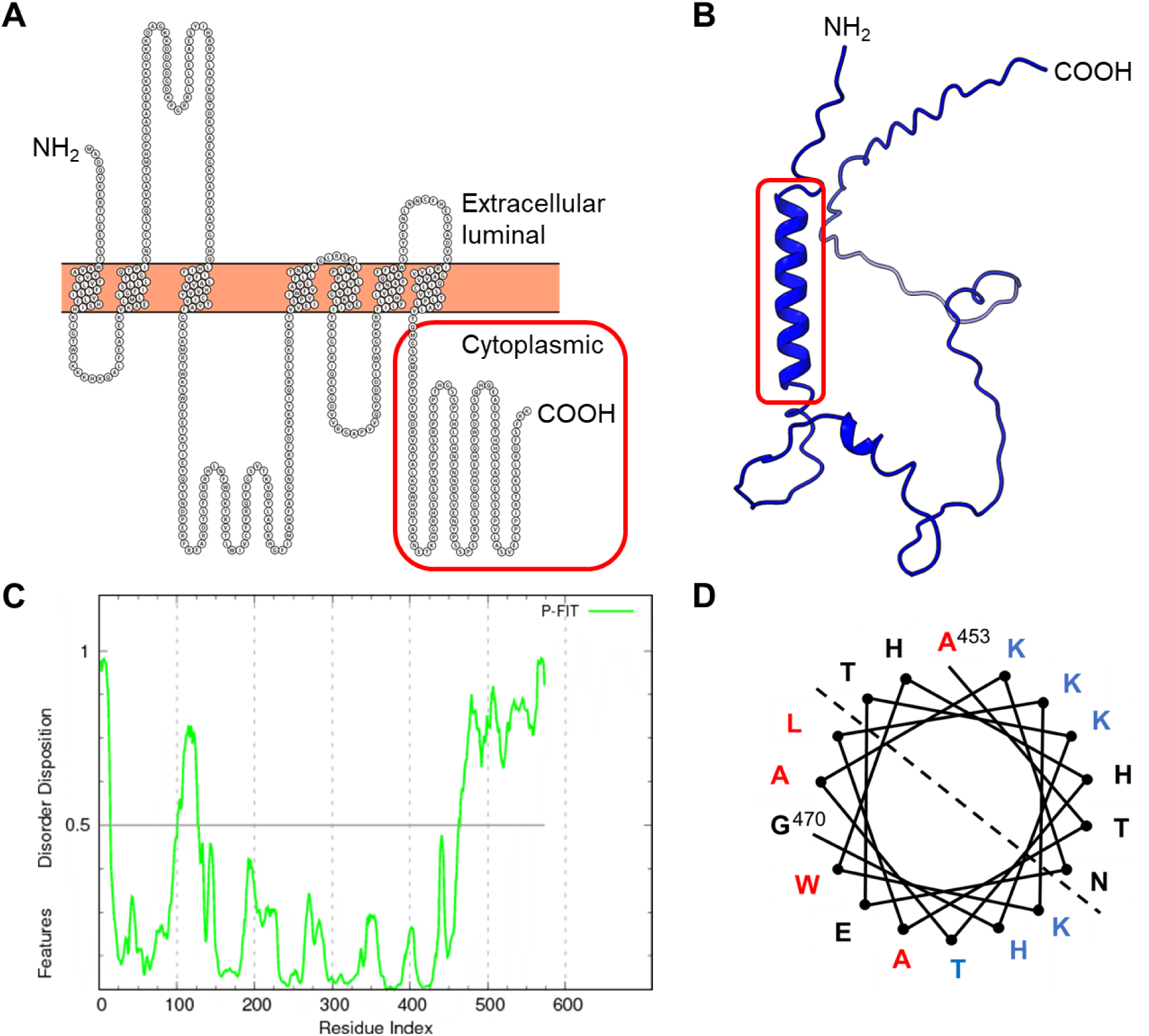
*In silico* analysis of the predicted MLO2^CT^ and its associated CAMBD. **A** Predicted membrane topology of MLO2. The amino acid sequence of the heptahelical MLO2 transmembrane protein was plotted with PROTTER (https://wlab.ethz.ch/protter/start/). The individual amino acids of MLO2 are presented as circles filled with the single-letter code for amino acids; the membrane is shown as an orange box. The amino terminus faces the extracellular/luminal side of the membrane; the cytoplasmic carboxyl terminus (starting from M^439^) is boxed in red. **B** Structure prediction of the MLO2^CT^ by AlphaFold. The carboxyl terminus (amino acids 439-573; corresponding to the boxed region in **A**) was subjected to structure prediction by AlphaFold. A predicted α-helical region between R^13^ (R^451^ according to the numbering of the full-length protein) and K^30^ (K^468^) is boxed in red. **C** Prediction of disordered protein regions in MLO2 by PONDR-FIT (http://original.disprot.org/pondr-fit.php). The plot shows the disorder disposition (y-axis) per amino acid position (x-axis). Regions with a score above 0.5 (indicated by the horizontal black line) are considered to be intrinsically disordered. **D** Helical wheel projection of the α-helical MLO2^CT^ region between A^15^ (A^453^ according to the numbering of the full-length protein) and G^32^ (G^470^). Individual residues are indicated corresponding to the single-letter code for amino acids. The dashed line separates one side of the helix with preferentially hydrophobic residues (red; bottom left) from another side of the helix with preferentially basic residues (blue; top right).

### Initial characterization of the MLO2^CT^-CAM2 interaction by a CAM overlay assay

The *A. thaliana* genome harbors seven *CAM* genes that encode for highly similar isoforms with a minimum of 96% identity between each other at the amino acid level. Three of the seven CAM isoforms (CAM2, CAM3, and CAM5) are even identical and a fourth isoform (CAM7) differs from these by only one amino acid (McCormack *et al*. 2005). We focused in the context of this work on CAM2 (At2g41110), which is a representative of the three identical isoforms.

To assess the putative binding of CAM2 to the CAMBD of MLO2, we first performed an *in vitro* CAM overlay assay using recombinant proteins. To this end, MLO2^CT^ (amino acids 439-573) of MLO2 was recombinantly expressed in *E. coli* as a fusion protein N-terminally tagged with glutathione *S*-transferase (GST). Both a WT version (MLO2^CT^) and a mutant variant harboring the L^18^R W^21^R (numbering according to the MLO2^CT^) double amino acid substitution (MLO2^CT-LW/RR^) within the MLO2 CAMBD were generated. This mutation is analogous to the one previously found to abolish CAM binding to barley and rice MLO proteins (Kim *et al*. 2002a; Kim *et al*. 2002b).

Furthermore, C-terminally hexahistidine-tagged CAM2 (CAM2-His_6_) was recombinantly expressed in *E. coli*, purified on nickel nitrilotriacetic acid (Ni-NTA) columns, and chemically linked to maleimide-activated horseradish peroxidase (HRP) *via* a stable thioether linkage to the reduced cysteine-39 residue of CAM2 (**Supplemental Figure 2**). For the actual overlay assay, lysates of *E. coli* strains expressing the GST-tagged MLO^CT^ variants were separated by sodium dodecyl sulfate polyacrylamide gel electrophoresis (SDS-PAGE) and transferred to a nitrocellulose membrane that was subsequently probed with the CAM2-His_6_-HRP conjugate. An empty vector control expressing GST only served as a negative control.

Immunoblot analysis with the α-GST antibody indicated expression of the full-length (∼41.5 kDa) GST-MLO2^CT^ and GST-MLO2^CT-LW/RR^ fusion proteins in *E. coli* and in both instances the presence of a cleavage product of lower molecular mass (∼35 kDa; **Figure 2**). The expression levels of the GST fusion proteins were similar to that of the GST only (empty vector; ∼29 kDa) control. The CAM overlay assay was performed on a separate membrane with CAM2-His_6_-HRP in the presence of 1 mM CaCl_2_, which revealed a strong signal for the full-length GST-MLO2^CT^ fusion protein, indicative of *in vitro* interaction between the two proteins. The low molecular mass cleavage product was also detectable with the α-GST antibody, suggesting that this protein fragment harbors the CAMBD of MLO2. No signal was detected for the GST-MLO2^CT-LW/RR^ fusion protein or the GST control in our conditions. Overlay of yet another membrane with CAM2-HRP in the presence of 5 mM of the calcium chelator ethylene glycol-bis(β-aminoethyl ether)-*N*,*N*,*N*′,*N*′-tetraacetic acid (EGTA) in addition to 1 mM CaCl_2_ completely prevented binding of CAM2-His_6_-HRP to either of the GST-MLO2^CT^ fusion proteins (**Figure 2**). In summary, results of the CAM overlay assay indicated Ca^2+^-dependent binding of CAM2-His_6_-HRP to the CAMBD of MLO2 (**Table 1**). This binding is prohibited by mutation of two amino acid residues (L^18^R and W^21^R, corresponding to L^456^R and W^459^R in full-length MLO2) that are in analogous positions within the CAMBD to those that were previously identified as being essential for the binding of CAM to barley and rice MLO proteins (Kim *et al*. 2002b; Kim *et al*. 2002a).

**Figure 2.**
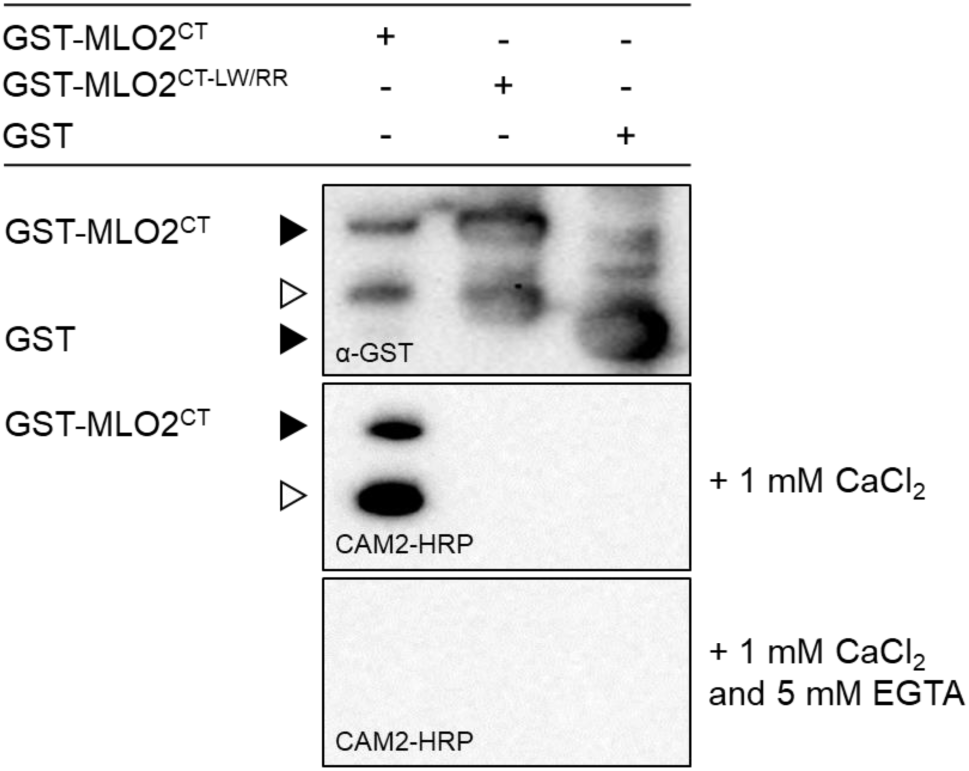
Initial characterization of the MLO2^CT^-CAM2 interaction by a CAM overlay assay. CAM2 overlay assay with recombinantly expressed HRP-labeled CAM2-His_6_ and GST-tagged MLO2^CT^ in the presence of either 1 mM CaCl_2_ (middle panel) or 1 mM CaCl_2_ plus 5 mM EGTA (lower panel). Protein loading was assessed by immunoblot analysis with an α-GST antibody (upper panel). Expected molecular masses of GST-MLO2^CT^ (∼41.5 kDa) and GST (∼29 kDa) are marked by a black triangle, a GST-MLO2^CT^ cleavage product (∼35 kDa) by a white triangle.

**Table 1.** Summary of data from the various MLO2-CAM2 interaction assays^a^.

|  | **MLO2** | **MLO2<sup>LW/RR</sup>** | **MLO2<sup>CT</sup>** | **MLO2<sup>CT-LW/RR</sup>** |
| --- | --- | --- | --- | --- |
| CAM overlay assay | n.t. | n.t. | +++ | - |
| GST pull-down assay | n.t. | n.t. | +++ | + |
| Y2H assays | n.t. | n.t. | - | - |
| Ura3-based yeast SUS | +++ | +++ | n.t. | n.t. |
| PLV-based yeast SUS | +++ | ++ | n.t. | n.t. |
| BiFC assay | n.t. | n.t. | +++ | +++ |
| LCI assay | +++ | + | +++ | + |
| TbID assay | +++ | + | n.t. | n.t. |

<sup>a</sup> compiled based on data shown in Figure 1-8. +++ strong interaction, ++ medium interaction, + weak interaction, - no interaction, n.t., not tested

### Analysis of site-directed MLO2^CT^ mutants *via* the CAM overlay assay

To find out whether the L^18^R and W^21^R amino acid substitutions within the CAMBD of MLO2 are the most effective mutations to abrogate CAM2 binding to the MLO2^CT^, we created a set of additional amino acid substitutions within the CAMBD and tested these *via* the above-described CAM overlay assay. We focused on six of the eight amino acid residues that are invariant between the barley Mlo and *A. thaliana* MLO2 CAMBDs (A^17^, L^18^, W^21^, A^25^, K^26^ and K^30^; **Supplemental File 1**) for site-directed mutagenesis. These residues represent amino acids for both the hydrophobic (A^17^, L^18^, W^21^ and A^25^) and basic (K^26^ and K^30^) side of the amphipathic α-helical CAMBD and, according to the helical wheel projection, reside in a conserved relative position within the Mlo and MLO2 CAMBDs (**Supplemental Figure 1**). In addition, we included H^31^, which is a further invariant amino acid among the highly conserved *A. thaliana* paralogs MLO2, MLO6 and MLO12. Hydrophobic amino acid residues were mutated to arginine (A^17^R, L^18^R, W^21^R and A^25^R), while hydrophilic ones were mutated to alanine (K^26^A, K^30^A and H^31^A). All variants were generated as N-terminally tagged GST fusion proteins by heterologous expression in *E. coli*.

Immunoblot analysis with the α-GST antibody indicated similar expression levels for all recombinant protein variants in *E. coli* and, as described above (**Figure 2**), the presence of a cleavage product of lower molecular mass that occurred in case of all variants (**Figure 3**). The CAM overlay assay in the presence of 1 mM CaCl_2_ revealed WT-like or possibly even stronger binding of CAM2-His_6_-HRP to the A^25^R, K^26^A, K^30^A and H^31^A MLO2^CT^ variants. Reduced binding of CAM2-His_6_-HRP was seen in case of the A^17^R variant, while no signal could be detected for the L^18^R and W^21^R single mutant variants, the L^18^R W^21^R double mutant variant, as well as the GST negative control. Signals were also absent for all the constructs in the presence of 5 mM EGTA, indicating the Ca^2+^-dependence of CAM binding (**Figure 3**). Taken together, this analysis revealed that the L^18^R and W^21^R amino acid substitutions as well as the L^18^R W^21^R double exchange are the most effective mutations to prevent the CAM binding to the CAMBD of MLO2 in the context of the CAM overlay assay.

**Figure 3.**
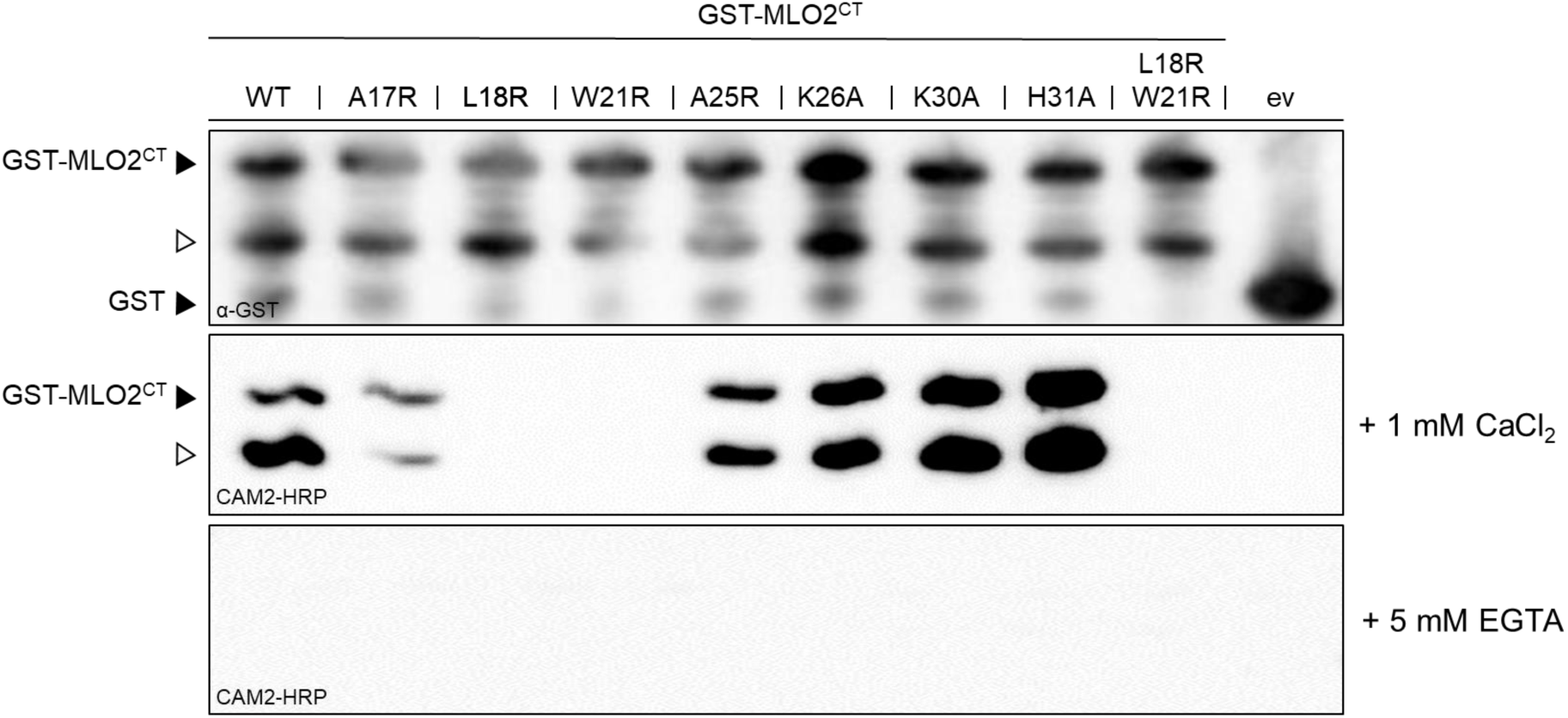
Analysis of site-directed MLO2^CT^ mutants *via* the CAM overlay assay. CAM2 overlay assay with recombinantly expressed HRP-labeled CAM2 and GST-tagged site-directed MLO2^CT^ mutant variants in the presence of either 1 mM CaCl_2_ (middle panel) or 1 mM CaCl_2_ plus 5 mM EGTA (lower panel). Protein loading was assessed by immunoblot analysis with an α-GST antibody (upper panel). Expected molecular masses of GST-MLO2^CT^ and GST are marked by a black triangle, a GST-MLO2^CT^ cleavage product by a white triangle. The assay was repeated twice with similar results. WT, wild-type version of the MLO2^CT^; ev, empty vector.

### Analysis of site-directed MLO2^CT^ mutants *via* a GST pull-down assay

We next aimed to validate the results of the CAM overlay assay with an independent *in vitro* experimental approach. To this end, we established a GST pull-down assay in which the GST-MLO2^CT^ was incubated with glutathione agarose beads to immobilize the fusion protein on a solid matrix. Purified hexahistidine-tagged CAM2 (CAM2-His_6_) was then added as a prey protein in the presence of 1 mM CaCl_2_, with or without 10 mM EGTA, and the mixtures were washed rigorously to remove unbound protein from the beads prior to the elution of the bound proteins from the glutathione agarose beads in SDS gel loading buffer and separation by SDS-PAGE. An initial experiment revealed strong Ca^2+^-dependent binding of CAM2-His_6_ to GST-MLO2^CT^ but strongly reduced binding of CAM2-His_6_ to the respective GST-MLO2^CT-LW/RR^ double mutant variant under these conditions (**Supplemental Figure 3**).

We extended the experiment by using *E. coli* cell homogenates of strains expressing the above-described set of GST-MLO2^CT^ variants as well as a L^18^R W^21^R A^25^R triple mutant variant and a version lacking the entire CAMBD (MLO2^CT-ΔBD^). Immunoblot analysis with α-GST and α-His antibodies indicated similar expression levels for all input samples. The GST-MLO2^CT^ samples showed, as described above for the CAM overlay assay (**Figure 2**), the presence of a cleavage product of lower molecular mass that occurred for all variants. In case of the pull-down samples in the presence of 1 mM CaCl_2_, the MLO2^CT^ wild type version resulted in a signal that was considerably stronger than that of the GST and GST-MLO2^CT-ΔBD^ negative controls. Wild-type-like or even stronger signals were seen for the A^17^R, A^25^R, K^26^A, K^30^A and H^31^A MLO2^CT^ variants. By contrast, we observed weak signals (comparable to the negative controls) for the L^18^R, W^21^R, L^18^R W^21^R and L^18^R W^21^R A^25^R MLO2^CT^ variants. Apart from faint background signals, the presence of 5 mM EGTA prevented the occurrence of signals for all tested constructs (**Figure 4**). Taken together, the results of the CAM overlay assay and the GST pull-down assay largely agree, except for the A^17^R variant, which yielded an inconsistent outcome in the two types of *in vitro* experiments. In both assays, the L^18^R W^21^R double mutant version lacked interaction with CAM2 (CAM overlay assay; **Figure 2** **and** **Figure 3; Table 1**) or showed a strong reduction in association (GST pull-down assay; **Figure 4; Table 1**). For the subsequent *in vivo* assays we, therefore, focused on the L^18^R W^21^R double mutant variants next to the respective MLO2 and MLO^CT^ WT versions.

**Figure 4.**
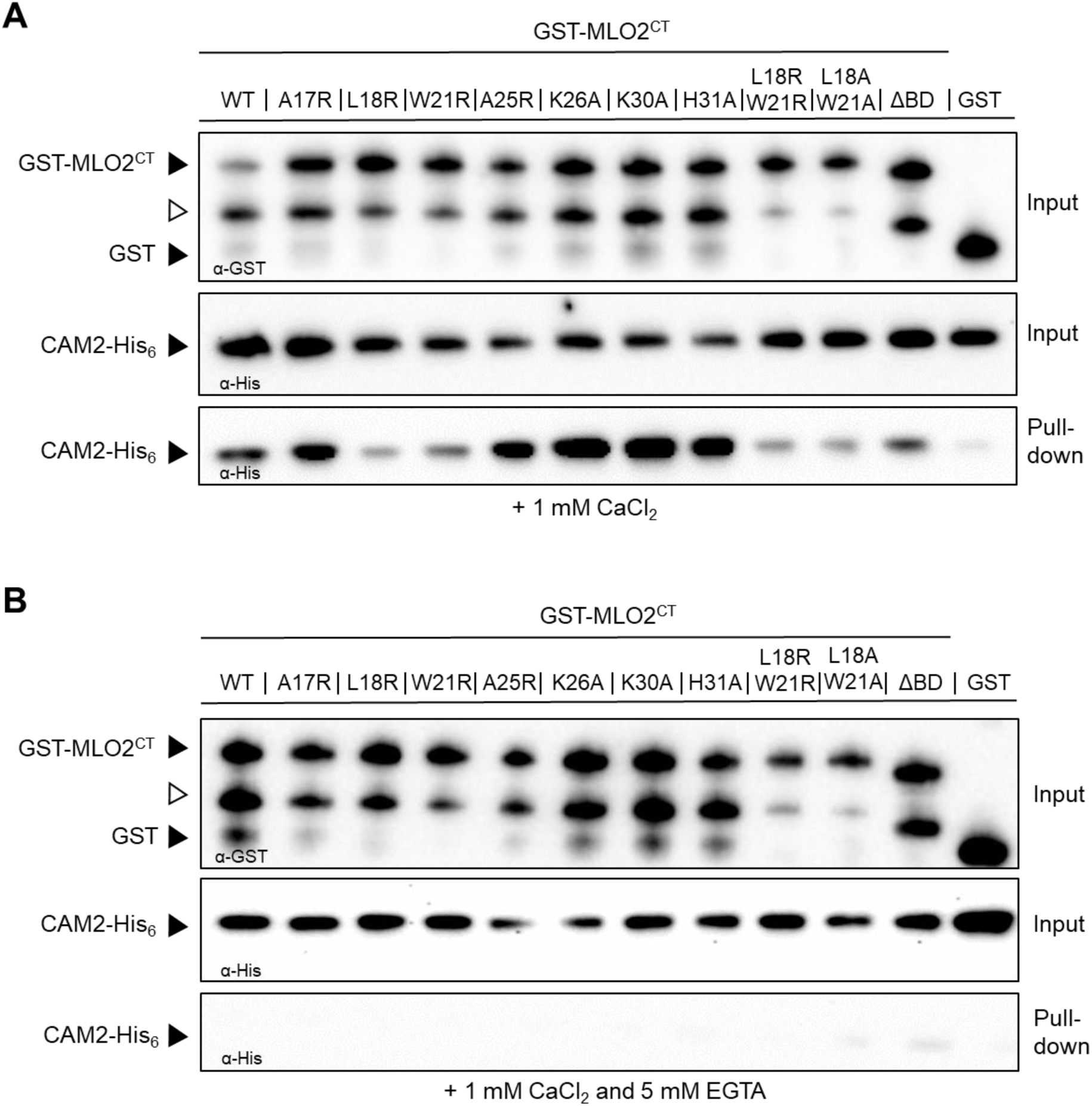
Analysis of site-directed MLO2^CT^ mutants *via* a GST pulldown assay. GST pulldown assay with recombinantly expressed CAM-His_6_ and GST-tagged site-directed MLO2^CT^ mutant variants in the presence of either 1 mM CaCl_2_ (**A**) or 1 mM CaCl_2_ plus 5 mM EGTA (**B**). Protein input was assessed by immunoblot analysis with α-GST (upper panel each) and α-His (middle panel each) antibodies. Presence of CAM-His_6_ was analyzed by immunoblot analysis with an α-His antibody (lower panel each). In the upper panel, expected molecular masses of GST-MLO2^CT^ and GST are marked by a black triangle, a GST-MLO2^CT^ cleavage product by a white triangle. The assay was repeated twice (in part with less mutant variants tested) with similar results. WT, wild-type version of the MLO2^CT^; ΔBD, version of the MLO2^CT^ lacking the entire CAMBD; i.e. amino acids A^17^ to H^31^ deleted; GST, GST tag alone (not fused to MLO2^CT^).

### Interaction between MLO2^CT^ or MLO2^CT-LW/RR^ and CAM2 in different yeast-based systems

In the following, we assessed the interaction between MLO2 and CAM2 *in vivo* using various yeast-based interaction assays. For the classical Y2H system, we employed two different commercially available and broadly used vector pairs that both enable N-terminal fusions of bait and prey proteins with the Gal4 transcription factor activation- and DNA-binding domains, respectively. While one pair comprises the low-copy vectors pDEST32 and pDEST22, the other consists of the high-copy vectors pGBKT7-GW and pGADT7-GW. Since the full-length MLO2 protein is membrane-localized and not able to enter the yeast nucleus, which is a prerequisite for interaction in the classical Y2H system, we focused on the MLO2^CT^ for the interaction studies with the Y2H method. We first tested the MLO2^CT^-CAM interaction with the low-copy vectors pDEST32 and pDEST22 in combination with the PJ69-4A yeast strain. Despite production of the proteins (**Supplemental Figure 4**), we did not observe evidence for the interaction between MLO2^CT^ or MLO2^CT-LW/RR^ and CAM2 in this setup, as indicated by the absence of any yeast growth on interaction-selective synthetic complete (SC) medium, which did not differ from the empty vector controls (**Figure 5A**). As we failed to detect any interaction with the pDEST32/pDEST22 vector system, we next moved to the pGBKT7-GW/pGADT7-GW high-copy vectors in combination with yeast strain AH109. In this setup, we analyzed both possible vector constellations for MLO2^CT^/MLO2^CT-LW/RR^ and CAM2. However, similar to the low-copy vector system, for none of the combinations tested we observed growth of the yeast colonies on interaction-selective SC medium (**Figure 5B**).

**Figure 5.**
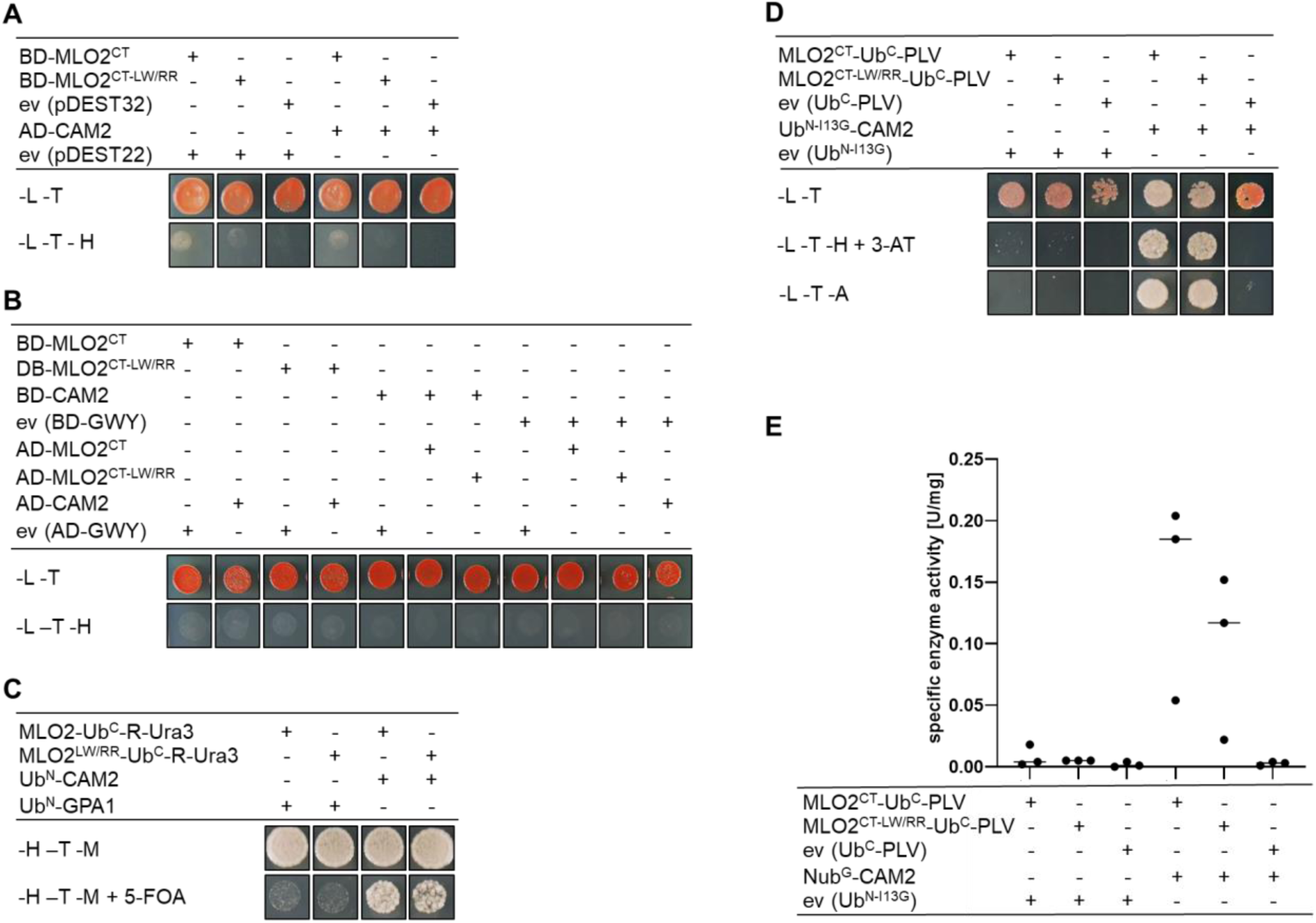
Interaction between MLO2^CT^ or MLO2^CT-LW/RR^ and CAM2 in different yeast-based systems. **A** Classical Y2H assay with the pGBKT7 (bait vector; Gal4 DNA-binding domain (BD); MLO2^CT^, MLO2^CT-LW/RR^ and empty vector) and pGADT7 (prey vector; Gal4 activation domain (AD); CAM2 and empty vector) vector system in *S. cerevisiae* strain AH109. Growth control was performed on SC medium lacking leucine (-L, selection for bait vector) and tryptophan (-T, selection for prey vector). Selection for interaction was performed on SC medium lacking leucine (-L), tryptophan (-T), and histidine (-H, selection for interaction). The assay was repeated twice with similar results. ev, empty vector. **B** Classical Y2H assay with the pDEST32 (bait vector; Gal4 DNA-binding domain (BD); MLO2^CT^ and MLO2^CT-LW/RR^ and empty vector) and pDEST22 (prey vector; Gal4 activation domain (AD); CAM2 and empty vector) vector system in *S. cerevisiae* strain PJ69-4A. Growth control was performed on SC medium lacking leucine (-L, selection for bait vector) and tryptophan (-T, selection for prey vector). Selection for interaction was performed on SC medium lacking leucine (-L), tryptophan (-T), and histidine (-H, selection for interaction). The assay was repeated twice with similar results. ev, empty vector. **C** Ura3-based yeast SUS with the pMet-GWY-Cub-R-Ura3 (bait vector; MLO2^CT^ and MLO2^CT-LW/RR^) and pCup-NuI-GWY-Cyc1 (prey vector; CAM2 and GPA1) vector system in *S. cerevisiae* strain JD53. Growth control was performed on SC medium lacking histidine (-H, selection for bait vector) and tryptophan (-T, selection for prey vector). Selection for interaction was performed on SC medium containing 0.7 g/L 5-FOA (+5-FOA, selection for interaction) and lacking methionine (-M, to allow for full promoter activity of the bait vector). All plates contained 500 µM methionine to reduce background growth due to a strong promoter activity of the bait vector. The assay was repeated twice with similar results. **D** PLV-based yeast SUS with the pMetOYC (bait vector; MLO2^CT^ and MLO2^CT-LW/RR^) and pNX32 (prey vector; CAM2) vector system in the *S. cerevisiae* strain THY.AP4. Growth control was performed on SC medium lacking leucine (-L, selection for bait vector) and tryptophan (-T, selection for prey vector). Selection for interaction was performed either on SC medium lacking leucine (-L), tryptophan (-T) and histidine (- H, selection for interaction) and in the presence of 10 mM 3-aminotriazole (+3-AT) or on SC medium lacking leucine (-L), tryptophan (-T) and adenine (-A, selection for interaction). All plates contained 500 µM methionine to reduce background growth due to a strong promoter activity of the bait vector. The assay was repeated twice with similar results. ev, empty vector. **E** Quantification of interaction strength in the PLV-based yeast SUS *via* a β-galactosidase reporter assay. Yeast cells were harvested from freshly grown cultures (OD = 1) and washed with sterile Z-buffer. Cells were disrupted by three freeze-and-thaw cycles, and the debris was separated by centrifugation. The protein concentration of the supernatant was determined. Aliquots of the supernatant, Z buffer, and the substrate o-nitrophenyl-β-D-galactopyranoside were mixed and incubated at 37 °C. The yellowing of the solution was monitored over time and stopped by the addition of Na2CO3 (final concentration 0.33 M). The extinction was measured at 420 nm, and the specific enzyme activity (U/mg) was calculated. Three independent biological replicates were performed and all data points are indicated. An ordinary one-way ANOVA followed by Tukeýs multiple comparison testing revealed no statistically significant differences between samples.

As we failed to detect any MLO2-CAM2 interaction in the classical Y2H, we next moved to the Ura3-based yeast SUS, which is suitable for analyzing the interaction of membrane proteins and, therefore, allows for the expression of full-length MLO2 and MLO2^LW/RR^ (Wittke *et al*. 1999). In this system, the bait protein (here: MLO2) is C-terminally fused to the C-terminal half of ubiquitin (Ub^C^) and the Ura3 (5-phosphate decarboxylase) reporter protein harboring an N-terminal destabilizing arginine (R) residue, while the prey protein (here: CAM2) is N-terminally tagged with the N-terminal half of ubiquitin (Ub^N^). Upon interaction between bait and prey proteins and reconstitution of ubiquitin, the pre-destabilized Ura3 reporter protein is proteolytically cleaved by ubiquitin-specific proteases, allowing for the growth of yeast cells on interaction-selective SC medium containing 5-fluoroorotic acid (5-FOA) (Wittke *et al*. 1999; Boeke *et al*. 1987). Using this yeast SUS setup, we noticed growth of yeast (strain JD53) transformants expressing full-length MLO2 and CAM2 on interaction-selective plates harboring 5-FOA. However, a similar level of yeast growth was seen in the case of the yeast transformants expressing the MLO2^LW/RR^ construct (**Figure 5B**). The heterotrimeric G-protein α-subunit GPA1 served as a prey negative control in this experiment.

Since interaction assays by means of the Ura3-based yeast SUS provide solely qualitative and no quantitative data and rest on a single reporter readout, we also opted for an alternative yeast SUS. The PLV-based yeast SUS depends on the interaction-dependent proteolytic release of an artificial multi-domain transcriptional activator comprised of a stabilizing protein A domain, a LexA DNA-binding domain and a VPS16 transactivation domain. Three different reporter genes (*His*, *Ade* and *LacZ*) can be activated by the liberated PLV transactivator upon interaction between the Ub^C^- and Ub^N^-tagged bait and prey proteins (Stagljar *et al*. 1998). We initially aimed at the expression of full-length MLO2 and MLO2^LW/RR^ in this yeast SUS. However, expression of these baits resulted in the constitutive activation of the reporter systems due to instability of the respective fusion proteins in our conditions. As an alternative, we deployed a modified version of the PLV-based yeast SUS in which cytosolic bait proteins are membrane-anchored *via* translational fusion with the yeast Ost4 membrane protein (Möckli *et al*. 2007). This yeast SUS variant enabled us to express MLO2^CT^ and MLO2^CT-LW/RR^ as C-terminal fusions with the Ub^C^ domain and the PLV transactivator in yeast strain THY.AP4 (**Supplemental Figure 4**). The CAM2 prey protein, on the other hand, was N-terminally fused with a Ub^N^ variant carrying an isoleucine to glycine substitution (I^13^G, Ub^N-I13G^) that reduces the affinity of Ub^N^ to Ub^C^ considerably, lowering the probability of false-positive interactions (Johnsson and Varshavsky 1994; Stagljar *et al*. 1998). Similar to the Ura3-based yeast SUS assay with MLO2 full-length proteins (see above; **Figure 5C**), this setup revealed interaction between MLO2^CT^ and CAM2 as well as MLO2^CT-LW/RR^ and CAM2 with no recognizable difference between the two bait proteins when considering yeast colony growth on selective media (**Figure 5D**). However, when measuring β-galactosidase activity as a quantitative readout of the *LacZ* reporter gene, we noticed that in each of three independent replicates enzymatic activity was lower for the yeast transformants expressing the MLO2^CT-LW/RR^ bait as compared to the corresponding yeast transformants expressing the MLO2^CT^ bait. Although this resulted in different median values for MLO2^CT^ (∼0.18 U/mg) and MLO2^CT-LW/RR^ (median ∼0.12 U/mg), the difference between the figures for the two bait variants was statistically not significant, likely due to the high experiment-to-experiment variation regarding absolute values in this assay (**Figure 5E**). In summary, while the Y2H assay failed to detect any MLO2^CT^-CAM2 interaction (**Figure 5A and B; Table 1**), the MLO2/MLO2^CT^-CAM2 interaction could be demonstrated by two different yeast SUS platforms. However, the presumed difference between the MLO2 WT version and the LW/RR mutant variant was, depending on the yeast system used, either not recognizable (**Figure 5C and D; Table 1**) or statistically not significant (**Figure 5E; Table 1**).

### Interaction between MLO2^CT^ or MLO2^CT-LW/RR^ and CAM2 visualized by a bimolecular fluorescence complementation (BiFC) assay

Next, we aimed to study the MLO^CT^-CAM2 interaction *in planta*. We first chose the BiFC system, which relies on bait and prey proteins tagged with the N- and C-terminal segments of the yellow fluorescent protein (YFP). Upon interaction of the bait and the prey protein, functional YFP may be reconstituted, yielding fluorescence upon appropriate excitation (Schütze *et al*. 2009; Walter *et al*. 2004).

We generated translational fusions of MLO2^CT^ and MLO2^CT-LW/RR^ with the C-terminal YFP segment (YFP^C^-MLO2^CT^- and YFP^C^-MLO2^CT-LW/RR^-, respectively) and CAM2, N-terminally tagged with the N-terminal YFP segment (YFP^N^-CAM2), and transiently co-expressed the YFP^C^-MLO2^CT^ / YFP^N^-CAM2 and YFP^C^-MLO2^CT-LW/RR^ / YFP^N^-CAM2 pairs in leaves of *N. benthamiana*. The MDL2-YFP^C^ / YFP^N^-CAM2 combination served as a negative control in this assay. MDL2 is a cytoplasmic protein (Gruner *et al*. 2021) assumed not to interact with CAM. At two days after infiltration of the agrobacteria, typically no or little fluorescence was detectable for any of the tested protein pairs. By contrast, at three days after infiltration of the agrobacteria, we observed clear fluorescence signals for the YFP^C^-MLO2^CT^ / YFP^N^-CAM2 and YFP^C^- MLO2^CT-LW/RR^ / YFP^N^-CAM2 pairs, while either no or weak fluorescence was seen for the negative control (MDL2-YFP^C^ / YFP^N^-CAM2) (**Figure 6**). However, we found no reproducible difference in fluorescence intensity between the combinations involving MLO2^CT^ and MLO2^CT-LW/RR^ (**Figure 6**). Thus, similar to the classical Y2H and the Ura3-based yeast SUS system (**Figure 5A-C**), the LW/RR double amino acid substitution in the MLO2^CT^ does not translate into a detectable difference in the BiFC interaction assay (**Table 1**).

**Figure 6.**
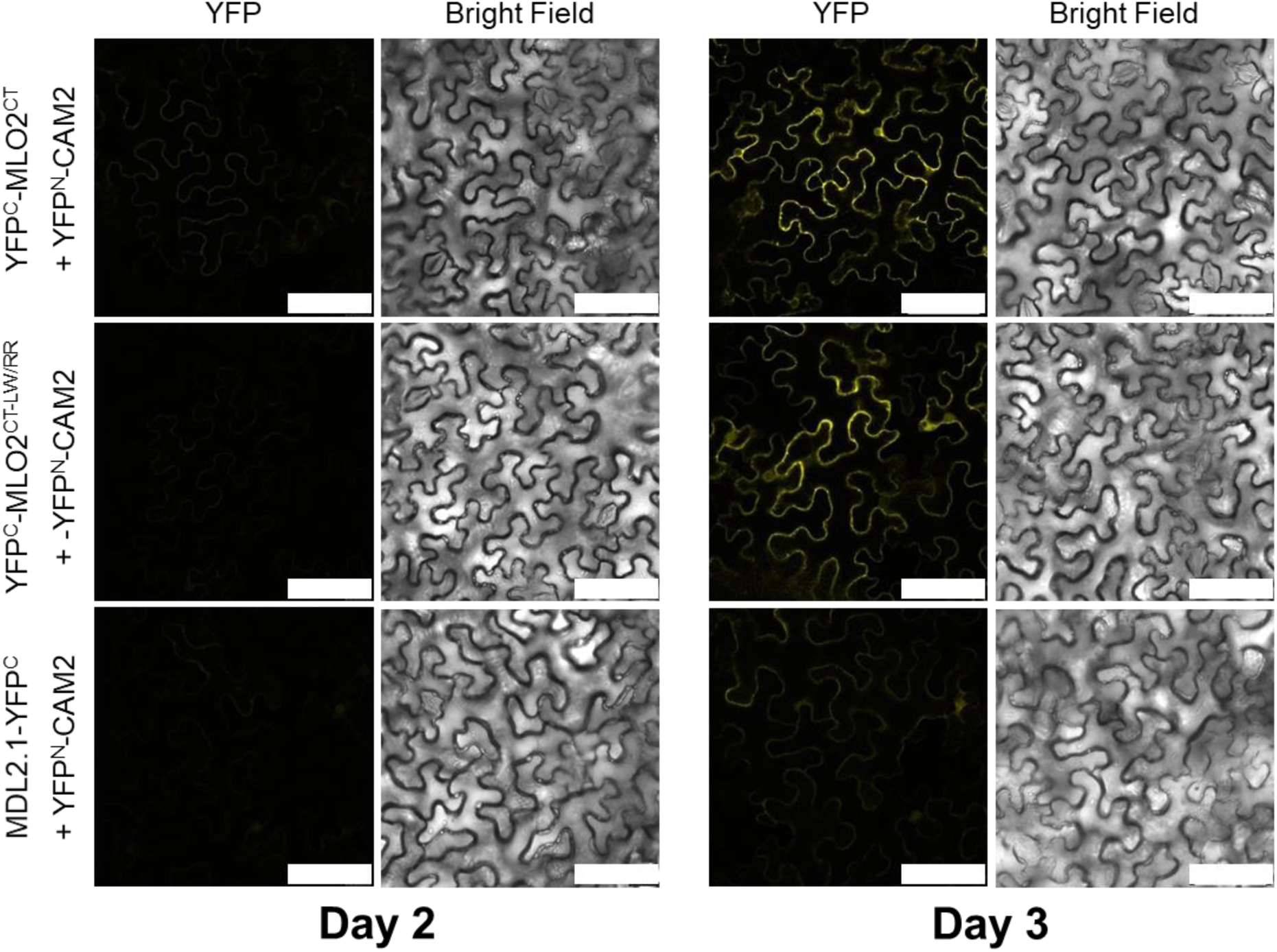
Interaction between MLO2^CT^ or MLO2^CT-LW/RR^ and CAM2 in a BiFC system. YFP^C^-MLO2^CT^ / YFP^N^-CAM2, YFP^C^-MLO2^CT-LW/RR^ / YFP^N^-CAM2, and MDL2.1-YFP^C^ / YFP^N^-CAM2 pairs were co-expressed in leaves of *N. benthamiana* by co-infiltration of *A. tumefaciens* strains harboring the respective plasmids. Leaves were analyzed by confocal laser scanning microscopy at two and three days after the infiltration of agrobacteria. Size bar, 75 µm. The experiment was repeated five times with similar results.

### Interaction between MLO2/MLO2^CT^ or MLO2 ^LW/RR^/MLO2^CT-LW/RR^ and CAM2 visualized by a Luciferase Complementation Imaging (LCI) assay

Similar to the BiFC assay, the LCI assay relies on the complementation of N- and C-terminal protein fragments (here from firefly luciferase, LUC). Reconstitution of the enzyme upon protein-protein interaction results in luciferase activity that can be measured in the presence of the substrate, luciferin (Chen *et al*. 2008). We first generated translational fusions of MLO2^CT^ and MLO2^CT-LW/RR^ with the N-terminal luciferase segment (LUC^N^-MLO2^CT^- and LUC^N^-MLO2^CT-LW/RR^-, respectively) and CAM2, N-terminally tagged with the C-terminal LUC segment (LUC^C^-CAM2), and transiently co-expressed the LUC^N^-MLO2^CT^ / LUC^C^-CAM2 and LUC^N^-MLO2^CT-LW/RR^ / LUC^C^-CAM2 pairs in leaves of *N. benthamiana*. As an additional control, an empty vector (LUC^N^) was used. We measured strong luciferase activity (median ∼46,000 units/mm^2^) in the case of the LUC^N^-MLO2^CT^ / LUC^C^-CAM2 combination and significantly reduced luciferase activity for the LUC^N^-MLO2^CT-LW/RR^ / LUC^C^-CAM2 pair (median ∼4,900 units/mm^2^). Comparatively low background luciferase activity (median ∼2,300 units/mm^2^) was seen when the LUC^N^ empty vector was co-infiltrated with LUC^C^-CAM2 (**Figure 7A** **and Supplemental Figure 5A**). In planta protein production was validated by immunoblot analysis (**Supplemental Figure 5B**). Taken together, this data set indicates reduced binding of CAM2 to MLO2^CT-LW/RR^ mutant variant in the context of the *in planta* LCI assay.

**Figure 7.**
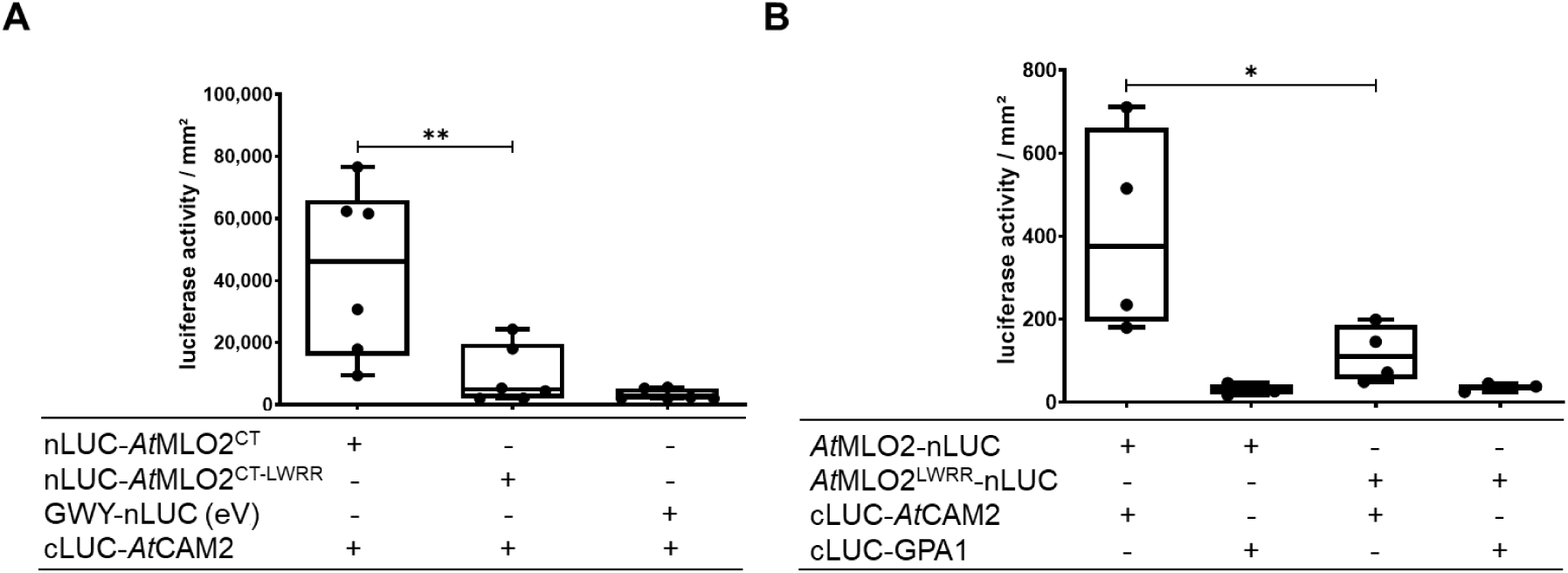
Interaction between MLO2^CT^ or MLO2^CT-LW/RR^ and CAM2 as well as MLO2 or MLO2^LW/RR^ and CAM2 in a LCI system. The indicated pairs including MLO2^CT^ (**A**) or full-length MLO2 (**B**) were co-expressed in leaves of *N. benthamiana* by co-infiltration of *A. tumefaciens* strains harboring the respective plasmids. **A** Leaves (one per biological replicate) were sprayed with luciferin at 3 days post infiltration of agrobacteria and luminescence quantified following dark incubation for 20 min. **B** For the experiment with full-length MLO2/MLO2^LW/RR^, leaf discs (12 per combination and biological replicate) were prepared at 3 days post infiltration, placed in 96-well plates, and luminescence was recorded after addition of luciferin and following dark incubation of 5 min. Five (**A**) or four (**B**) independent biological replicates were performed. Asterisks indicate a statistically significant difference between MLO2^CT^ or MLO2 in comparison to the respective LW/RR double mutant variant according to one-way ANOVA followed by Tukey’s multiple comparison test (* *p*<0.05, ** *p*<0.01). See **Supplemental Figure 4** for a representation based on relative light units for this assay.

We next wondered whether this result could be recapitulated in the context of the full-length MLO2 protein. To this end, we generated LCI constructs in which full-length MLO2 WT and a respective LW/RR (L^456^R W^459^R) mutant variant were C-terminally tagged with the N-terminal luciferase fragment (MLO2-LUC^N^) and co-expressed these transiently in *N. benthamiana* with CAM2, N-terminally tagged with the C-terminal luciferase fragment (LUC^C^-CAM2). In this set of experiments, the *A. thaliana* heterotrimeric G-protein α-subunit, GPA1, N-terminally tagged with the C-terminal luciferase fragment (LUC^C^-GPA1), served as a negative control. In comparison to the negative control combinations (MLO2-LUC^N^ / LUC^C^-GPA1 and MLO2^LW/RR^-LUC^N^ / LUC^C^-GPA1; median luciferase activity ∼35 units/mm^2^ each), we measured marked luciferase activity for the MLO2-LUC^N^ / LUC^C^-CAM2 pair (∼375 units/mm^2^; **Figure 7B**). This measured value is substantially lower than the figure obtained in the context of the LUC^N^-MLO2^CT^ / LUC^C^-CAM2 combination (median ∼46,000 units/mm^2^; **Figure 7A**), which is likely due to different expression levels of MLO2^CT^ and full-length MLO2 and/or due to methodological differences in the assays (see Materials and Methods for details). Notably, similar to the experiment with the MLO^CT^, the MLO2^LW/RR^-LUC^N^ / LUC^C^-CAM2 pair yielded significantly lower luciferase activity (median ∼110 unit/mm^2^; **Figure 7B**), indicative of reduced CAM2 binding to MLO2.

When normalized against the respective negative controls (empty vector in the case of MLO2^CT^ and LUC^C^-GPA1 in the case of full-length MLO2), the relative light units were similar for the WT and LW/RR variants in the two assays (**Supplemental Figure 6**). In summary, both N-terminally tagged MLO2^CT^ and C-terminally tagged MLO2 full-length protein interact with CAM2 in the LCI assay, and the respective LW/RR mutant variants exhibit in each case reduced interaction (**Table 1**).

### Interaction between MLO2 or MLO2^LW/RR^ and CAM2 visualized by a proximity-dependent biotin labeling assay

We finally tested the MLO2-CAM2 interaction by proximity-dependent biotin labeling. To this end, MLO2 and MLO2^LW/RR^ fusion proteins with TurboID (TbID) were transiently co-expressed with epitope-labeled CAM2 in *N. benthamiana*. TbID is an improved biotin ligase that uses ATP to convert biotin into biotinol-5′–AMP, a reactive intermediate that covalently labels lysine residues of nearby proteins. Subsequent streptavidin immunoprecipitation enriches for biotin-labeled target proteins, which can be further analyzed, e.g. by immunoblot analysis (Yang *et al*. 2021).

Dexamethasone-inducible expression of MLO2-TbID or MLO2^LW/RR^-TbID in combination with CAM2, N-terminally labeled with a hemagglutinin (HA) tag (HA-CAM2), in the presence of 250 µM biotin resulted in a wide spectrum of biotinylated proteins, covering a broad molecular mass range. Although the overall pattern was similar, the intensity of biotin labeling appeared to be stronger at 24 h as compared to 6 h after biotin application. We did not observe any obvious difference in the labeling pattern between the expression of MLO2-TbID and MLO2^LW/RR^-TbID (**Figure 8A**).

**Figure 8.**
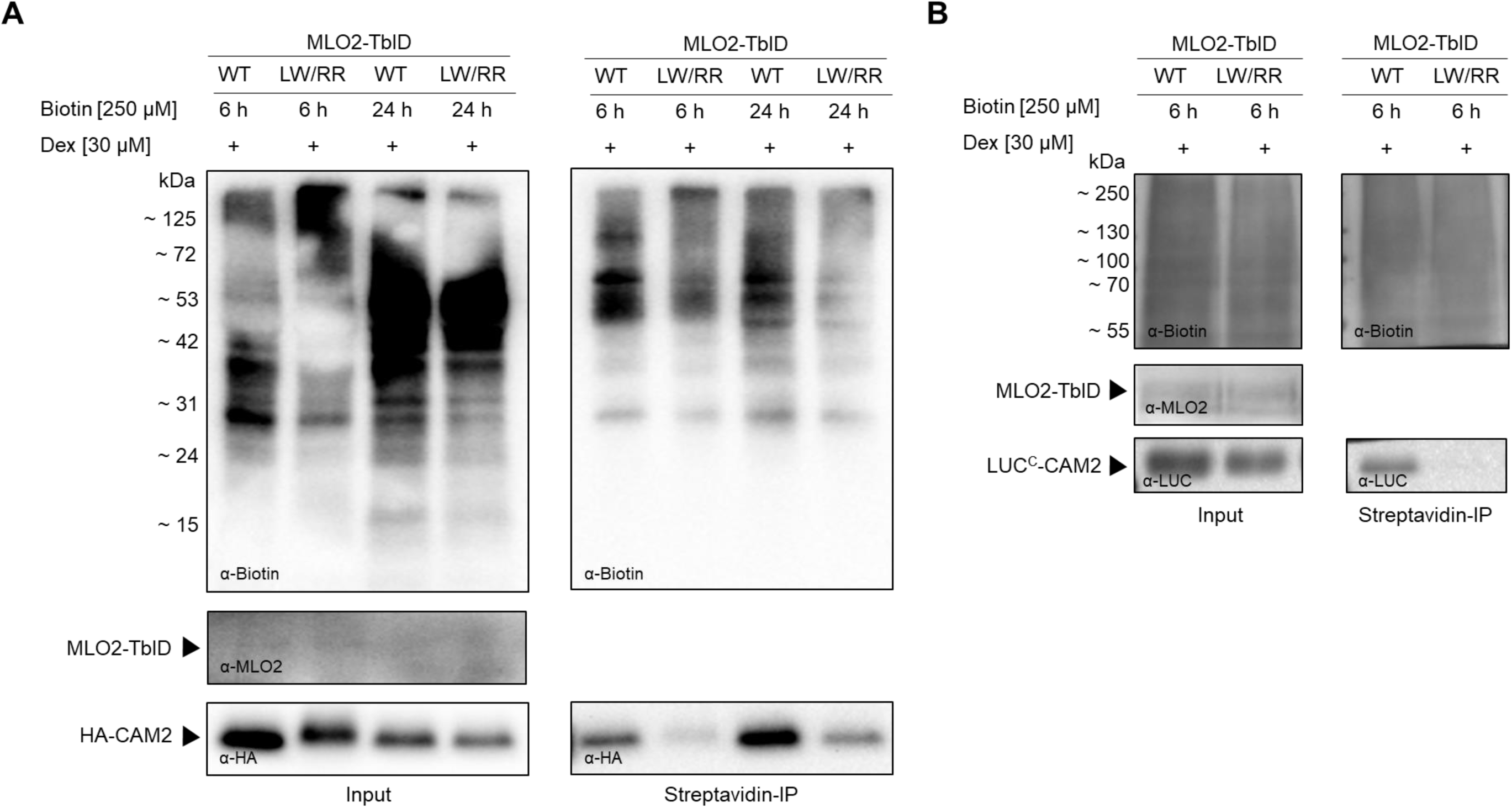
Interaction between MLO2^CT^ or MLO2^CT-LW/RR^ and CAM2 by a proximity-dependent biotin labeling assay. MLO2-TbID / HA-CAM2 or MLO2^LW/RR^-TbID / HA-CAM2 (**A**) or MLO2-TbID / LUC-CAM2 or MLO2^LW/RR^-TbID / LUC^C^-CAM2 (**B**) were co-expressed in leaves of *N. benthamiana* in the presence of 250 µM biotin by co-infiltration of *A. tumefaciens* strains harboring the respective plasmids. Expression of MLO2-TbID or MLO2^LW/RR^-TbID was induced by the addition of 30 µM dexamethasone (Dex); biotin solution (250 µM) was infiltrated 24 h later and proteins extracted either 6 h or 24 h thereafter, as indicated above the immunoblots. Biotinylated proteins were immunoprecipitated with streptavidin beads. Proximity-dependent biotin labeling of total protein extracts prior (Input; left panels) and after (Streptavidin-IP; right panels) immunoprecipitation was analyzed by immunoblot with an α-biotin antibody (upper panel). Input expression of MLO2-TbID was validated by immunoblot with an α-MLO2 antiserum (left; middle panel). CAM2 input expression (left; lower panel) and biotin labeling (right; lower panel) were analyzed by immunoblots with an α-HA (**A**) or α-LUC (**B**) antibody, respectively.

After immunoprecipitation of the biotinylated proteins with streptavidin beads, we recovered a similar spectrum of biotin-labeled proteins, although proteins of lower molecular mass appeared to be somewhat underrepresented. Immunoblot analysis of the immunoprecipitated sample with an α-HA antibody for the detection of HA-CAM2 revealed marked levels of this protein upon expression of MLO2-TbID at 6 h after biotin application, indicating the intracellular presence of HA-CAM2 in the vicinity of MLO2-TbID. We detected an even stronger accumulation of HA-CAM2 at 24 h after biotin application, consistent with an assumed increased biotinylation of this target protein over time. In comparison to MLO2-TbID, we noticed reduced band intensities for HA-CAM in the immunoprecipitated samples upon expression of MLO2^LW/RR^-TbID, both at 6 h and 24 h after biotin application, suggesting a reduced association of MLO2^LW/RR^-TbID and HA-CAM under these conditions (**Figure 8A**). To validate this outcome, we repeated the experiment using a different epitope tag (LUC^C^) N-terminally fused to CAM2, focusing on 6 h biotin application, which yielded the more pronounced difference between MLO2-TbID and MLO2^LW/RR^-TbID in the first trial.

Similar to the experiment with HA-CAM2 (**Figure 8A**), co-expression of LUC^C^-CAM2 with MLO2^LW/RR^-TbID yielded substantially lower levels of biotinylation than co-expression of LUC-CAM2 with MLO2-TbID (**Figure 8B**). Thus, TbID-mediated biotin proximity labeling is suitable to visualize the MLO2-CAM2 interaction and sensitive enough to discriminate WT and the LW/RR mutant variant (**Table 1**).

## Discussion

We here studied the interaction between *A. thaliana* MLO2 (or its C-terminus harboring the CAMBD) and CAM2 with seven different experimental approaches. In each type of assay, we deployed both the wild-type version of the CAMBD (either in the context of the MLO2^CT^ or the full-length MLO2 protein) and at least the respective LW/RR double mutant. Except for the classical Y2H approach, each of the methods indicated association of CAM2 with either the MLO2 full-length protein or the MLO2^CT^ (**Table 1**). Previously, interaction between MLO proteins and either CAM or CML proteins was seen in several cases with a variety of methods (Zhu *et al*. 2021; Kim *et al*. 2002a; Kim *et al*. 2002b; Kim *et al*. 2014; Yu *et al*. 2019; Bhat *et al*. 2005). A comprehensive Y2H study revealed that the C-termini of all 15 *A. thaliana* MLO proteins can interact with at least one CML (Zhu *et al*. 2021). Our results using MLO2 further strengthen the notion that the interaction of MLO proteins with CAM/CMLs is a common feature of MLO proteins that likely contributes to their *in vivo* functionality.

The data further validate the C-terminal CAMBD as the primary contact site between MLO and CAM/CML proteins, although the residual association of CAM2 with MLO2 LW/RR mutant variants could point at a contribution by additional domains of the protein (see below). Although all results of this study were obtained with CAM2, we believe that due to the high sequence conservation among the seven *A. thaliana* CAM isoforms with a minimum of 96% sequence identity, the outcomes of our interaction assays are likely to be representative for all CAMs encoded by the Arabidopsis genome.

We tested site-directed mutants of seven amino acids that are conserved between the CAMBD of MLO2 and barley Mlo, or between the CAMBDs of MLO2, MLO6 and MLO12 (A^17^, L^18^, W^21^, A^25^, K^26^, K^30^ and H^31^; **Supplemental Figure 1**) in a CAM overlay assay. This revealed, similar to a previous study with barley Mlo (Kim *et al*. 2002b), protein variants with unaltered (A^25^R, K^26^A), reduced (A^17^R, L^18^R, W^21^R) and enhanced (K^30^A, H^31^A) *in vitro* CAM binding capacity. Especially the latter feature is remarkable since it suggests that at least barley Mlo and *A. thaliana* MLO2 proteins did not evolve their maximal CAM binding affinity, at least as judged from the *in vitro* assays. This may indicate that CAM binding to MLO proteins is a fine-tuned and balanced process, highlighting its putative physiological relevance in the context of MLO function. Results of a recent study indicate that calcium-dependent CAM association with the MLO CAMBD might be required for autoinhibition of MLÓs calcium channel activity (Gao *et al*. 2022). It is conceivable that the extent of this negative feedback activity may differ dependent on the particular MLO paralog and its respective cellular and physiological context.

We focused in our study in particular on the LW/RR double amino acid substitution within MLO2 CAMBD and its ability to interact with CAM– either in the context of the MLO2 full-length protein or its cytoplasmic C-terminus (MLO2^CT^). We subjected this constellation to seven different protein-protein interaction assays: (1) CAM overlay assay (**Figure 2 and 3**), (2) GST pull-down assay (**Figure 4**), (3) two versions of the classical Y2H assay (**Figure 5A and B**), (4) two variants (Ura3- and PLV-based) of the yeast SUS assay (**Figure 5C-E**), (5) BiFC assay (**Figure 6**), (6) LCI assay (**Figure 7**) and (7) proximity-dependent biotin labeling assay (**Figure 8**). In the case of five of the mentioned experimental approaches (CAM overlay, GST pull-down, classical Y2H, PLV-based yeast SUS, and LCI), MLO2^CT^ and its corresponding MLO2^CT-LW/RR^ mutant variant were offered as potential interaction partners for CAM2. Similarly, for another four techniques (Ura3-based yeast SUS, BiFC, LCI and biotin labeling), the MLO2 full-length protein was deployed (note that in the case of the yeast SUS and *in planta* LCI assay both MLO2^CT^ and full-length MLO2 were tested). Two of the mentioned methods (CAM overlay and GST pull-down) are *in vitro* test systems, three (Y2H as well as Ura3- and PLV-based yeast SUS) rely on yeast, and another three (BiFC, LCI, and biotin labeling) are *in planta* assays. The majority of the procedures tested (CAM overlay, GST pull-down, PLV-based yeast SUS, LCI, and biotin labeling) revealed either a qualitative or quantitative difference in the interaction between the MLO2/MLO2^CT^ LW/RR double mutant and CAM in comparison to the respective WT versions (**Figure 2**, **Figure 3**, **Figure 4**, **Figure 5C and D**, **Figure 7** **and** **Figure 8**). These differences in the strength of CAM binding are unlikely to be the result of lower expression levels of the MLO2^LW/RR^ and MLO2^CT-LW/RR^ mutant variants in relation to the respective WT versions as we controlled in most assays (apart from BiFC) for equal protein expression levels by immunoblot analysis. Our data, thus, corroborate a critical role of the highly conserved amino acid residues in the CAMBD of MLO proteins.

Exceptions from the differential outcome between MLO2/MLO2^CT^ WT and mutant versions were the classical Y2H approach, which failed to detect any interaction between MLO2^CT^ and CAM (**Figure 5A and B**), as well as the Ura3-based yeast SUS and the BiFC assay, which did not discriminate between the MLO2 WT and LW/RR variants (**Figure 5B** **and** **Figure 6**). The BiFC system is known to be prone to false-positive results due to the high tendency of self-association of the two halves of the fluorescent proteins, which, once formed, constitute an irreversible complex, thereby stabilizing interactions between any fused interaction partners (Xing *et al*. 2016; Miller *et al*. 2015). While this feature can be an advantage for the detection of transient protein-protein interactions, it is usually considered a disadvantage since it may result in the formation of artificial protein complexes due to random protein-protein contacts. Accordingly, mutant variants were highly recommended to be included as essential controls in BiFC experiments (Kudla and Bock 2016). In comparison to the BiFC assay, the outcome of the Ura3-based yeast SUS experiment was unexpected, as a similar assay with another *A. thaliana* MLO family member, MLO1, previously revealed reduced interaction with its respective LW/RR mutant variant (Kim *et al*. 2002b). Likewise unexpected was the failure to detect any interaction between MLO2^CT^ and CAM2 in the Y2H since a previous study found interactions between *A. thaliana* MLO family members (including MLO2^CT^) and CAM-like proteins (CMLs) using the pGBKT7/pGADT7-based Y2H also deployed in our study (**Figure 5B**). While canonical CAMs harbor four calcium-binding EF hands, CML proteins have a variable number of one to six EF hands and, accordingly, typically differ in the total number of amino acids from classical CAMs. The interaction partners of the MLO2 carboxyl-terminus identified in the study of Zhu and co-workers (Zhu *et al*. 2021), CML9 and CML18, harbor four EF hands each and have a similar number of amino acids as CAM2 tested in our work (151 and 161 as compared to 149 amino acids). However, these proteins share only 50% (CML9) and 45% (CML19) sequence identity and 69% (both proteins) sequence similarity with CAM2, which may explain the differential outcome in the Y2H assays performed before (Zhu *et al*. 2021) and in the present study (**Figure 5A and B**).

It is noteworthy that the CAM overlay assay, similar to previous findings with barley and rice MLO (Kim *et al*. 2002a; Kim *et al*. 2002b), revealed a seemingly complete absence of the interaction between the MLO2^CT-LW/RR^ mutant and CAM2, even at possibly unphysiologically high calcium concentrations (**Figure 2** **and** **Figure 3**). By contrast, most of the tested *in vivo* approaches (PLV-based yeast SUS, LCI and biotin labeling) rather point to a reduced level of association between the MLO2 LW/RR mutant variant and CAM2 (**Figure 5D and E**, **Figure 7** **and** **Figure 8**). While experimental details may account for this discrepancy between the different methods, there might also be biological explanations. One possibility is that the mutated CAMBD indeed exhibits residual binding affinity for CAM/CML proteins under *in vivo* conditions. Another option is that further cytoplasmic domains of MLO2, such as its large second cytoplasmic loop (Devoto *et al*. 1999; Kusch *et al*. 2016; Devoto *et al*. 2003), affect the MLO2-CAM2 interaction *in planta*, e.g. by stabilizing an initial association of the two binding partners. In addition or alternatively, further proteins present in the yeast and plant cells of the respective *in vivo* assays could modulate the interaction. It needs, however, to be considered that all protein-protein interaction assays performed in the context of this study were based on unphysiologically high protein concentrations due to overexpression. Therefore, the residual binding of CAM2 to the mutated MLO2 CAMBD in the *in vivo* assays could simply represent an overexpression artefact.

Our transient gene expression experiments in *N. benthamiana* revealed *in vivo* biotinylation of CAM2 by the TbID biotin ligase C-terminally fused to MLO2 (**Figure 8**). While this approach was used in the context of the present work to probe the MLO2-CAM2 interaction, it could be deployed in future studies to identify novel interaction partners of MLO proteins. Apart from CAM (Kim *et al*. 2002a; Kim *et al*. 2002b; Kim *et al*. 2014; Zhu *et al*. 2021) cyclic nucleotide gated channels (CGNCs; Meng *et al*. 2020) and exocyst EXO70 subunits (Huebbers *et al*. 2022), no other plant proteins have been reported to date to associate *in planta* with MLO proteins. Being integral membrane proteins, the identification of protein interaction partners is notoriously difficult for MLO proteins. The TbID approach promises to capture physiologically relevant *in vivo* protein-protein interactions, possibly also in different cell types and in different physiological contexts (Zhang *et al*. 2020; Mair *et al*. 2019; Arora *et al*. 2020; Yang *et al*. 2021). To this end, future experiments should involve the expression of functionally validated MLO-TbID fusion proteins in stable transgenic lines, ideally driven by the corresponding native *MLO* promoter.

## Materials and Methods

### *In silico* predictions

The membrane topology of *A. thaliana* MLO2 (At2g11310; https://www.uniprot.org/uniprot/Q9SXB6) was determined and drawn using PROTTER (https://wlab.ethz.ch/protter/start/). We used the predicted cytoplasmic C-terminal region of MLO2 (MLO2^CT^) for further *in silico* analyses. Analogous to other MLO proteins (Piffanelli *et al*. 1999; Panstruga 2005; Kusch *et al*. 2016; Devoto *et al*. 2003), the MLO2^CT^ region starts after the last predicted transmembrane domain with a methionine residue (M^439^) and comprises amino acids 439-573, i.e., 135 residues in total. The numbering of the amino acids within this study refers to M^439^ in the full-length protein as M^1^ in the MLO2^CT^. The PONDR-FIT tool (http://original.disprot.org/pondr-fit.php; Xue *et al*. 2010), a meta-predictor of intrinsically disordered proteins, was employed to predict disordered regions within the MLO2 protein. The AlphaFold (Jumper *et al*. 2021) prediction of three-dimensional structure of the MLO2^CT^ was run at https://colab.research.google.com/github/sokrypton/ColabFold/blob/main/AlphaFold2<u>. ipynb?pli=1#scrollTo=kOblAo-xetgx. The rank 1 model was chosen for visualization with ChimeraX (Pettersen *et al*. 2021). Helical wheel projections were calculated by pepwheel (</u>https://www.bioinformatics.nl/cgi-bin/emboss/pepwheel<u>) and wheel graphs drawn manually. All tools were used using default parameters.</u>

### Cloning of expression constructs

The *MLO2^CT^* coding sequence was originally inserted as an *Nco*I/*Eco*RI DNA fragment into *E. coli* vector pGEX-2TK (GE Healthcare Life Sciences, Chalfont St. Giles, U.K.) for the inducible high-level expression of GST-MLO2^CT^ fusion protein. Site directed mutagenesis of *MLO2^CT^* was performed by Gibson assembly (Gibson *et al*. 2009) based on suitable PCR fragments generated with Phusion^®^ high-fidelity DNA polymerase (NEB GmbH, Frankfurt, Germany). The *CAM2* coding sequence was inserted as an *Nco*I/*Xho*I DNA fragment into modified pET28a vector that lacks the N-terminal His_6_ tag (previously designated pETλHIS; Campe *et al*. 2016) for the inducible high-level expression of CAM2-His_6_ fusion protein.

Constructs for the Y2H and yeast SUS assays were generated by Gateway^®^ cloning. *MLO^CT^* and *MLO2^CT-LW/RR^* were shuttled into pDEST32 (Invitrogen - Thermo Fisher Scientific, Waltham, MA, USA), pGBKT7 and pGADT7 (Clontech, now Takara Bio, San Jose, CA, USA), as well as in pMETOYC-Dest (Xing *et al*. 2016), but in the latter without a stop codon. Full-length *MLO2* and *MLO2^LW/RR^* genes (lacking a stop codon) were transferred by Gateway^®^ LR reactions from pDONR entry clones into pMET-GWY-Cub-R-Ura3-Cyc1 (Deslandes *et al*. 2003; Wittke *et al*. 1999). Arabidopsis *CAM2* was shuttled by Gateway^®^ LR reactions into pDEST22 (Thermo Fisher Scientific), pGBKT7 and pGADT7 (Clontech), pCup-NuIGWY-Cyc1 (Wittke *et al*. 1999; Deslandes *et al*. 2003) and pNX32-Dest (Obrdlik *et al*. 2004).

Plasmid constructs used for the BiFC assay (pUBQ-cYFP-MLO2^CT^, pUBQ-cYFP-MLO2^CT-LW/RR^) were also generated by Gateway^®^ cloning. Inserts were moved by Gateway^®^ LR reactions from pDONR entry clones into destination vectors pUBN-YFP^C^ (Grefen *et al*. 2010), pE-SPYNE and pE-SPYCE (Walter *et al*. 2004) for BiFC assays.

For LCI assays, inserts were shuttled by Gateway^®^ LR recombination into either pAMPAT-LUC^N^ (used for MLO2^CT^ and MLO2^CT-LW/RR^) and pAMPAT-LUC^C^ (used for CAM2) -both for N-terminal tagging with LUC fragments (Gruner *et al*. 2021), or into pCAMBIA1300-N-LUC-GWY (for C-terminal tagging with LUC^N^; used for MLO2^CT^ and MLO2^CT-LW/RR^) and pCAMBIA1300-GWY-C-LUC (for N-terminal tagging with LUC^C^; used for CAM2) (Chen *et al*. 2008).

The dexamethasone-inducible MLO2-TbID construct is based on expression vector pB7m34GW (Karimi *et al*. 2005) and was generated by MultiSite Gateway™ technology to insert the dexamethasone-inducible pOp6/LhGR promoter system (Samalova *et al*. 2005) in front of the *MLO2* coding sequence, C-terminally fused to *TbID* (Branon *et al*. 2018) followed by a His_6_ epitope tag (MLO2-TbID-His_6_). To create the pOp6/LhGR-containing entry clone, the pOp6/LhGR module from vector pOp/LhGR was combined with the backbone of vector p1R4_G1090:XVE by Gibson assembly to replace the XVE module. The resulting donor plasmid, pG1090::LHGR/pOP6, has P4-P1r Gateway^®^ recombination sites. The *TbID-His_6_* coding sequence present in vector TurboID-His_6__pET21a (https://www.addgene.org/107177/; Branon *et al*. 2018) was recloned into pDONR P2r-P3 (Invitrogen - Thermo Fisher Scientific) and a stop codon introduced after the His_6_ tag. Finally, entry clones harboring the pOp6/LhGR promoter system, the *MLO2* coding sequence (in pDONR221 (Invitrogen – Thermo Fisher Scientific), lacking a stop codon) and the *TbID-His_6_* fragment were jointly recombined into vector pB7m34GW by MultiSite Gateway™ recombination. The corresponding MLO2^LW/RR^ construct was created by site-directed mutagenesis on the basis of Gibson assembly (Gibson *et al*. 2009) as described above. The plasmid for the *in planta* expression of HA-CAM2 was made by Gateway^®^-based transfer of the *CAM2* coding sequence into pEarleyGate201 (Earley *et al*. 2006).

### Generation of *E. coli* lysates

For the generation of bacterial lysates, 2 mL of an overnight culture of *E. coli* ROSETTA^TM^ (DE3) pLysS or BL21 (DE3) cells containing the appropriate expression constructs was transferred into 200 mL LB medium with appropriate antibiotics. The culture was incubated at 37 °C while shaking at 220 revolutions per minute (rpm) until OD600 reached 0.6-0.8. Protein expression was induced by addition of 1 mM isopropyl β-D-1-thiogalactopyranoside (IPTG) and incubation was continued at 28 °C for 3 h at 220 rpm. The cells were harvested by centrifugation at 3,130 x *g* for 15 min. The pellet was then dissolved in 8 mL lysis buffer (25 mM HEPES (pH 7.5), 300 mM NaCl, 10% (v/v) glycerol, 5 mM imidazole) and incubated at 4 °C while gently shaking for 30 min. The suspension was sonicated on ice for 2 min and centrifuged at 3,130 x *g* and 4 °C for 50 min. The bacterial lysate was either stored in 2 mL aliquots at -20 °C or immediately used for further analysis.

### Affinity purification of recombinant hexahistidine-labeled CAM2

The Protino^®^ Ni-NTA column (Macherey-Nagel, Düren, Germany) was used for affinity chromatography of recombinant CAM2-His_6_ from *E. coli* lysate. First, the column was equilibrated with 10 mL lysis buffer (see above) according to the manufacturer’s instructions. The bacterial lysate (see above) was loaded onto the column and then washed with 30 mL of wash buffer (25 mM HEPES (pH 7.5), 300 mM NaCl, 10% (v/v) glycerol, 20 mM imidazole). Finally, the hexahistidine-tagged protein was eluded with 5 mL elution buffer 25 mM HEPES (pH 7.5), 300 mM NaCl, 10% (v/v) glycerol, 300 mM imidazole) in five fractions of 500 μL each. A small sample (approx. 50 μL) of flow through was collected after each step for further analysis by SDS-PAGE (see below). Following elution of His_6_-CAM2, the buffer was exchanged with 1x phosphate-buffered saline (PBS; 137 mM NaCl, 2.7 mM KCl, 10 mM Na2HPO4, 1.8 mM KH2PO4 (pH ∼7.3-7.4)) using a PD-10 desalting column (GE Healthcare) according to the manufacturer’s instructions. The protein concentration was calculated by using a Nanodrop^TM^ 2000c spectrophotometer (Thermo Fisher Scientific) to measure absorbance at 280 nm.

### SDS-PAGE and immunoblot analysis

For SDS-PAGE, the Mini-PROTEAN^®^ Tetra cell (Bio-Rad, Hercules, CA, USA) was used. Bis-Tris-polyacrylamide gels were prepared consisting of 12% resolving gels and 4% stacking gels. Gels were run at room temperature in either 1x MES (50 mM 2-(*N*-morpholino)ethanesulfonic acid (MES), 50 mM Tris, 1 mM EDTA, 0.1% SDS; pH 7.3) or Laemmli running buffer (25 mM Tris, 250 mM glycine, 0.1% SDS) at 175 V for 45 min. As a molecular mass marker, 2.5 μL of PiNK/BlueStar Prestained Protein Marker (NIPPON Genetics EUROPE GmbH, Düren, Germany) was used per gel lane. After electrophoresis, gels were either directly stained with Instant Blue™ (Biozol, Eching, Germany), or the proteins were transferred onto a nitrocellulose membrane using Mini Trans-Blot^®^ cell (Bio-Rad, Hercules, CA, USA). The transfer was performed in 1x transfer buffer at 250 mA for 1 h at 4 °C under constant stirring. The membrane was blocked in 5% skim milk (w/v) in Tris-buffered saline with Tween-20 (20 mM Tris-HCl pH7.5, 150 mM NaCl, 0.1% Tween-20; TBST) for 1 h while gently shaking. Afterwards, the membrane was washed in 1x TBST three times for 5 min each and then incubated with the appropriate primary antibody at 4 °C overnight. The membrane was washed in 1x TBST three times for 5 min, before incubating with the secondary antibody for 1 h at room temperature. After washing again three times with TBST for 15 min, the presence of horseradish peroxidase (HRP) coupled to the secondary antibody was detected by addition of either SuperSignal West Pico substrate for strong bands or SuperSignal West Femto solution (Thermo Fisher Scientific, Waltham, MA, USA) for faint bands by using ChemiDoc™ XRS+ (Bio-Rad, Hercules, CA, USA) and ImageLab™ software. Finally, the membrane was washed with ddH2O and stained in Ponceau solution. After drying for several minutes, pictures were taken of the stained membrane to verify equal loading of proteins.

### Antibodies

For immunoblot analyses, the following commercially available primary antibodies were used: rabbit α-GST (Cell Signaling Technology, Danvers, MA, USA; used in 1:1,000 dilution), mouse α-His (Cell Signaling Technology; used in 1:1,000 dilution), rat α-HA (Hoffmann-La Roche AG, Basel, Switzerland; used in 1:1,000 dilution), goat α-luciferase (Sigma Aldrich, St. Louis, MO, USA; used in 1:1,000 dilution), rabbit α-Gal4 BD (Santa Cruz Biotechnology, Dallas, TX, USA; used in 1:1,000 dilution), rabbit α-Gal4 AD (Santa Cruz Biotechnology; used in 1:1,000 dilution), and goat α-biotin-HRP (Cell Signaling Technology; used in 1:2,000 dilution). In addition, we deployed a polyclonal rabbit α-MLO2 antiserum raised against the recombinantly expressed MLO2 carboxyl-terminus (used in 1:500 dilution) as well as a custom-made polyclonal rabbit α-LexA antiserum (used in 1:5,000 dilution) raised against the C-terminal 15 amino acids of PLV (Harty and Römisch 2013; kindly provided by Prof. Dr. Karin Römisch). As secondary antibodies, α-goat-HRP (Santa Cruz Biotechnology), α-mouse HRP (Thermo Fisher Scientific) α-rabbit-HRP (Cell Signaling Technology) and α-rat-HRP (Sigma Aldrich) were used as appropriate (all used in 1:2,000 dilution). Antibody dilutions were made in 5% (w/v) bovine serum albumin (α-GST, α-His, α-HA, α-biotin-HRP, α-MLO2) in TBST or 5% (w/v) milk (α-Gal4 BD, α-Gal4 AD, α-LUC and all secondary antibodies) in TBST.

### Labeling of CAM2 with HRP

For conjugation of HRP to CAM2-His_6_, 50 μL of 10 mM Tris(2-carboxyethyl)phosphine (TCEP) was added to 1 mL (corresponding to ∼1 mg) of purified CAM2-His_6_ and incubated at room temperature for 2 h to reduce all cysteine residues present in the protein. Thereafter, the TCEP was removed using a PD-10 desalting column (GE Healthcare) according to the manufacturer’s instructions. The reduced CAM2-His_6_ protein was then mixed with 1 mg EZ-Link™ Maleimide Activated HRP (Thermo Fisher Scientific) in a molar ratio of 1:1 and incubated overnight at room temperature. The next day, glycerol was added to the CAM2-HRP complex to reach a final concentration of 20 % (v/v). Successful linkage was validated by SDS-PAGE and subsequent Coomassie staining of the gel using Instant Blue™ (Biozol, Eching, Germany) (**Supplemental Figure 1**).

### CAM overlay assay

*E. coli* lysates of strains expressing the various constructs were mixed with 6x SDS loading buffer and samples boiled at 95 °C for 5 min before loading onto three separate Bis-Tris-polyacrylamide gels. After gel separation, proteins were transferred to nitrocellulose membranes. The membranes intended for the overlay assay were rinsed with 1x TBST and then blocked in 7% (w/v) milk in TBST overnight at 4 °C. After washing three times with 1x TBST, the membranes were subsequently equilibrated for 1 h in 20 mL overlay buffer (50 mM imidazole-HCl (pH 7.5), 150 mM NaCl) which additionally contained either 1 mM CaCl_2_ or 5 mM EGTA (also present in all subsequently used buffers). Next, the membranes were incubated at room temperature for 1 h in 20 mL overlay buffer with 0.1 % gelatin (w/v) and 1:1,000 diluted CAM2-HRP (∼20 µg – see above). Afterwards, the membranes were washed five times for 5 minutes in wash buffer 1 (1x TBST, 0.1 % Tween (v/v), 50 mM imidazole-HCl, (pH 7.5), 2 (20 mM Tris-HCl (pH 7.5), 0.5 % Tween (v/v), 50 mM imidazole-HCl (pH 7.5), 0.5 M KCl) and 3 (20 mM Tris-HCl (pH 7.5), 0.1 % Tween (v/v), 0.5 M KCl). Chemiluminescence was detected by addition of either SuperSignal West Pico substrate for strong bands or SuperSignal West Femto solution (Thermo Fisher Scientific) for faint bands by using ChemiDoc™ XRS+ (Bio-Rad, Hercules, CA, USA) and ImageLab™ software. Presence of equal protein amounts was validated by immunoblot analysis with an α-GST antibody.

### GST pull-down assay

For the pull-down assay with GST-tagged proteins, Protino^®^ Glutathione Agarose 4B (Macherey-Nagel) was used. For each reaction, 100 μL of thoroughly mixed slurry was washed with 1x PBS according to the manufacturer’s instructions and then resuspended in 100 μL of 1x PBS. The input of *E. coli* lysate was adjusted to 1.9 mL of the lowest concentrated lysate using the previously calculated relative protein amount. All following steps were performed on ice to prevent protein degradation. Glutathione sepharose beads (100 μL) and the cell lysate were mixed in a 2 mL reaction tube. The samples were filled up to 2 mL with 1x PBS and incubated for at least 1 h at 4 °C while rotating end-over-end at 25 rpm. Afterwards, the beads were collected by centrifugation at 500 x *g* for 5 min at 4 °C and then washed four times with 1 mL 1x PBS. After resuspension of the samples in 500 μL binding buffer (140 mM NaCl, 2.7 mM KCl, 10 mM Na2HPO4, 1.8 mM KH2PO4), two different reactions were prepared for each construct: 250 μL of bead suspension were mixed with 0.5 μL 1 M CaCl_2_ and 20 μg purified CAM2-His_6_. In addition, 20 μL of 250 mM EGTA (pH 8.0) was added to one half of the samples. The volume was filled up to 500 μL with binding buffer and then incubated at 4 °C for 1 h while rotating end-over-end at 25 rpm. Finally, the beads were washed five times with 1 mL wash buffer (400 mM NaCl, 2.7 mM KCl, 10 mM Na2HPO4, 1.8 mM KH2PO4, supplemented with either 1 mM CaCl_2_ or 5 mM EGTA) and resolved in 6x SDS loading buffer (12 % SDS (w/v), 9 mM bromophenol blue, 47% glycerol, 60 mM Tris-HCl (pH 6.8), 0,58 M DTT). After boiling for 10 min at 95 °C and shortly spinning the beads down, immunoblot analysis with α-GST and α-His antibodies was performed.

### Yeast-based interaction assays

Yeast cells were transformed with a modified LiAc protocol (Gietz and Woods 2002). A liquid overnight culture was grown at 30 °C and 250 rpm in YPD, SC-Leu or SC-His, depending on the yeast strain used. The main culture was set to OD 0.2 and was incubated until it reached an OD of 0.8-1. Cells were harvested by centrifugation at 1,500 x *g* for 5 min and washed with 30 mL sterile water. Afterwards, the cells were resuspended in 1 mL 1x TE (10 mM Tris-HCl pH 7.5, 1 mM EDTA)/1x LiAc (100 mM). Then, 1 µg of DNA and 50 µg of high-quality sheared salmon sperm DNA (Invitrogen-Thermo Fisher Scientific) as carrier DNA were added to a 50 µl aliquot of competent cells. Next, 300 µL of sterile 40% PEG-4000/1x LiAc/1x TE were combined with the mixture of cells and gently mixed. The cell suspension was incubated at 30 °C for 30 min and then shifted to 42 °C for a heat shock. The heat treatment lasted for 10 min for *S. cerevisiae* strains PJ69-4A (James *et al*. 1996) and AH109 (Clontech), used for the classical Y2H assays, and for 1 h for strains THY.AP4 (Grefen *et al*. 2009) and JD53 (Dohmen *et al*. 1995) used for the yeast SUS experiments. Transformed cells were plated on SC medium lacking appropriate amino acids (Formedium, Norfolk, UK) as selection markers (**Supplemental Table 1**) and grown for at least two days at 30 °C.

Expression of bait and prey constructs in yeast was verified *via* immunoblot with α-Gal4 BD, α-Gal4 AD (Santa Cruz Biotechnology) or α-LexA (Harty and Römisch 2013) antibodies. Protein extraction was performed with a modified protocol of the Dohlman lab for trichloroacetic acid (TCA) yeast whole cell extracts (adapted from (Cox *et al*. 1997); https://www.med.unc.edu/pharm/dohlmanlab/resources/lab-methods/tca/). In short, a 10 mL culture (OD 1) was harvested and resuspended in 300 µl of TCA buffer (10 mM Tris-HCl, pH 8.0; 10% trichloroacetic acid; 25 mM NH4OAc; 1 mM Na2 EDTA). Glass beads were added for cell disruption in 5 x 1 min bursts on a vortex. The cell lysate was transferred to a new tube, and the beads were washed with 100 µL TCA buffer and added to the new tube. The supernatant was removed after centrifugation for 10 min at 16,000 x *g* at 4 °C and resuspended in 150 µL resuspension solution (0.1 M Tris-HCl, pH 11; 3% SDS). The samples were boiled for 5 min and cell debris was separated by centrifugation for 30 sec at 16,000 x *g*. From the supernatant, 120 µL were transferred to a new tube and an aliquot thereof used for protein concentration measurements. Expression of the bait construct in the Ura3-based yeast SUS was validated by a growth assay on SC-His-Ura plates (not shown).

Drop tests to examine for protein-protein interactions were performed by harvesting and washing cells from overnight cultures of the respective strains, carrying bait and prey constructs, and diluting these to OD 1. A 10-fold dilution series was performed, and 4 µL of each dilution was dropped on suitable SC plates lacking specific amino acids or containing 3-AT (Y2H) or 5-FOA (Ura3-based yeast SUS; **Supplemental Table 1**). Plates were incubated for 2 to 4 days, and representative pictures were taken for documentation. The LacZ reporter assay was performed with a modified protocol of Clonetech. A freshly grown 10 ml (start OD 0.2) was grown to OD 1 and harvested by centrifugation (3,400 x *g* for 1 min). Cells were washed once with 1 mL sterile 4 °C-cold Z buffer (60 mM Na2HPO4 2 H2O, 40 mM NaH2PO4 H2O, 10 mM KCl, 1 mM MgSO4 7 H2O; pH 7.0) and then resuspended in 650 µL of Z buffer. To disrupt the yeast cells, three freeze and thaw cycles were accomplished in liquid nitrogen. After the addition of 50 µL 0.1% SDS and 50 µL chloroform, the solution was mixed for 1 min. The cell debris and lysate were separated by centrifugation at 10.000 x *g* for 10 min (4 °C). Of the supernatant, 600 µL were transferred to a new tube and a Bradford assay (Bradford 1976) was performed to determine protein concentration. To start the enzymatic reaction, 800 µL prewarmed (37 °C) oNPG-solution (1 mg/mL ortho-nitrophenyl-β-galactoside in Z buffer) was mixed with 200 µL yeast protein extract, which was diluted to the lowest protein concentration. The yellowing of the solution was monitored over time during the incubation time (at 37°C) and was stopped by adding 0.5 mL 1 M Na2CO3 before saturation. The extinction at 420 nm (E420) was measured and put into the following equation to calculate the specific enzymatic activity: [U/mg] = (E420 x V) / (ε x d x v x t x P), with V = volume of the reaction (1,500 μL), ε = extinction coefficient of o-nitrophenol (4,500 M^-1^ cm^-1^), d = thickness of the cuvette (1 cm), v = volume of yeast extract (200 μL) and t = reaction time.

### Bimolecular fluorescence complementation (BiFC) assay

For BiFC assays, constructs on the basis of vectors pUBN-YFP^C^ (Grefen *et al*. 2010) and pE-SPYNE and pE-SPYCE (Walter *et al*. 2004) were used. Leaves of 4-6 week-old *N. benthamiana* plants grown in short-day conditions (10 h light, 23 °C, 80-90% relative humidity, 80-100 µmol s^-1^ m^-2^ light intensity) were infiltrated with *A. tumefaciens* strains carrying the genes of interest that were tagged with either the N- or C-terminal part of yellow fluorescence protein (YFP) as follows: pUBQ::cYFP-MLO2^CT^ (pUBN-YFP^C^), pUBQ::cYFP-MLO2^CT-LW/RR^ (pUBN-YFP^C^), p35S::nYFP-CAM2 (pE-SPYNE), p35S::MDL2.1-cYFP (pE-SPYCE). In addition, an *A. tumefaciens* strain (GV2260) carrying the viral gene silencing suppressor p19 was co-infiltrated. After recovery for either two or three days in long-day conditions (16 h light, 20 °C, 60-65% relative humidity, 105-120 µmol s^-1^ m^-2^ light intensity), three leaf discs representing every tested interaction were stamped out, analyzed by confocal laser scanning microscopy (see below) and then frozen in liquid nitrogen for protein extraction and subsequent immunoblot analysis.

### Confocal laser scanning microscopy

Leaf discs punched from *Agrobacterium*-infiltrated *N. benthamiana* leaves (see section 2.2.12) were placed on a glass slide in ddH2O and then analyzed with a Leica TCS SP8 LIGHTNING Confocal Microscope (Leica Camera AG, Wetzlar, Germany) using the HC PL APO CS2 20x0.75 IMM objective. The fluorescence signal of YFP was analyzed by exciting at 514 nm with an argon ion laser and measuring emission at 520-550 nm.

### LCI assay

Leaves of 4-6-week-old *N. benthamiana* plants grown short-day conditions (10 h light, 23 °C, 80-90% relative humidity, 80-100 µmol s^-1^ m^-2^ light intensity) conditions were infiltrated with either *A. tumefaciens* strain GV3101 (pMP90RK) (for MLO2^CT^ constructs) or *A. tumefaciens* strain AGL1 (for MLO2 full-length constructs) carrying the genes of interest that were tagged with either the N- or C-terminal part of firefly luciferase. In addition, an *A. tumefaciens* strain (GV2260) carrying the viral gene silencing suppressor p19 was co-infiltrated. For testing MLO2^CT^ constructs, expression vectors pAMPAT-LUC^N^ and pAMPAT-LUC^C^ (Gruner *et al*. 2021) were used and the following constructs generated by Gateway^®^ LR recombination: p35S::LUC^N^-MLO2^CT^ (pAMPAT-LUC^N^), p35S::LUC^N^-MLO2^CT-LW/RR^ (pAMPAT-LUC^N^), p35S::LUC^C^-CAM2 (pAMPAT-LUC^C^) and p35S::LUC^N^ (pAMPAT-LUC^N^). For testing full-length MLO2 constructs, pCAMBIA1300-C-LUC-GWY and pCAMBIA1300-GWY-N-LUC (Chen *et al*. 2008) were used and the following constructs generated by Gateway^®^ LR recombination: p35S::MLO2-LUC^N^ (pCAMBIA1300-GWY-N-LUC), p35S::MLO2^LW/RR^-LUC^N^ (pCAMBIA1300-GWY-N-LUC), p35S::LUC^C^-CAM2 (pCAMBIA1300-C-LUC-GWY) and p35S::LUC^C^-GPA1 (pCAMBIA1300-C-LUC-GWY).

After recovery for three days in long-day conditions (16 h light, 20 °C, 60-65% relative humidity, 105-120 µmol s^-1^ m^-2^ light intensity), the leaves were sprayed with 1 mM D-luciferin (PerkinElmer, Rodgau, Germany) solution containing 0.01 % Tween-20 (v/v) and incubated in the dark for 20 min. Chemiluminescence was detected by using ChemiDoc™ XRS+ (Bio-Rad, Hercules, CA, USA) and ImageLab™ software. Three leaf discs were taken close from each agroinfiltration site for protein extraction and immunoblot analysis to validate protein expression. Alternatively, for full-length MLO2/MLO2^LW/RR^, twelve leaf discs per combination of constructs were taken close from agroinfiltration sites of a minimum of three different leaves (max. four discs/leaf). The leaf discs were placed in individual wells of a white 96-well plate containing 100 µL 10 mM MgCl_2_ per well. Prior measurement, the liquid was replaced by 100 µL of freshly prepared 10 mM MgCl_2_ containing 1 mM D-Luciferin. Following a dark incubation of 5 min, luminescence was recorded for 1 sec/well in a CENTRO luminometer (Berthold Technologies, Bad Wildbad, Germany). All twelve leaf discs per construct were pooled for protein extraction and immunoblot analysis to validate protein expression. Chemiluminescence values are given as relative light units per measured leaf area (RLU/mm^2^).

### Proximity-dependent biotin labeling assay

*Agrobacterium tumefaciens* GV3101 (pMP90RK) strains carrying the constructs pB7m34GW-MLO2, pB7m34GW-MLO2^LW/RR^, pEarleyGate-HA-CAM2 or pAMPAT-LUC^C^-CAM2 were mixed in respective combinations with *A. tumefaciens* strain GV2260, carrying the viral gene silencing suppressor p19, and infiltrated into leaves of 4-6-week-old *N. benthamiana* plants grown in short-day conditions (10 h light, 23°C, 80-90% relative humidity, 80-100 µmol s^-1^ m^-2^ light intensity). After two days of recovery in long-day conditions (16 h light, 20 °C, 60-65% relative humidity, 105-120 µmol s^-1^ m^-2^ light intensity), the leaves were sprayed with 30 µM dexamethasone (Dex) solution and incubated for another 24 h. Then, biotin solution (250 µM) was infiltrated into the leaves and samples were taken after 6 h and 24 h. A simple protein extraction from *N. benthamiana* tissue was performed with subsequent buffer exchange via P10 desalting columns, and all biotinylated proteins were bound by Pierce^TM^ streptavidin agarose beads (Thermo Fisher Scientific). To this end, 40 µL of the beads were washed three times (2,500 x *g*, 1 min) with 500 µL 8 M urea binding buffer (8 M urea, 200 mM NaCl, 100 mM Tris-HCl pH 8.0). After the last washing step, the beads were resuspended in 100 µL urea binding buffer and 40 µg of protein extract was added. The volume was adjusted to 40 µL with urea binding buffer and the samples were incubated over night at 25 rpm at room temperature. The samples were washed 5 times with 1 mL urea binding buffer and used for SDS PAGE and immunoblot analysis with α-biotin, α-MLO2, α-HA and α-LUC antibodies. The appropriate volume of SDS loading buffer was added to the immunoprecipitated protein samples and then boiled for 10 min at 95 °C. A share of the total protein extract was used for analysis of the input sample.

### Phenolic total protein extraction

Plant tissue was homogenized with metal beads by freezing the tubes in liquid nitrogen. For the whole extraction, every step was performed on ice, with pre-chilled solutions and with centrifuges set at 4 °C. The leaf powder was washed twice with 900 µL 100% acetone and centrifuged at 20.800 x *g* for 5 min. Afterwards, the pellet was dissolved in 900 µL 10% (w/v) TCA in acetone and the samples were exposed to ultrasound in an ice bath for 10 min. The samples were centrifuged again and washed 900 µL 10% (w/v) TCA in acetone, 900 µL 10% (w/v) TCA in H2O and 900 µL 80% (v/v) acetone. The pellet was resuspended in 300 µL freshly prepared dense SDS buffer (100 mM Tris-HCl pH 8.0, 30% (w/v) sucrose; 2% (w/v) SDS, 5% (v/v) β-mercaptoethanol) at room temperature and 300 µL phenol was added. The solution was mixed rigorously, and the phases were separated by centrifugation at room temperature for 20 min. Of the upper phase, 180 µL was mixed with 900 µL of 100 mM ammonium acetate in methanol. After an incubation for 1 h at -20 °C, the precipitate was collected by centrifugation, and the pellet was washed once with 900 µL 100 mM ammonium acetate in methanol and twice with 900 µL 80% (v/v) acetone. The dry pellet was resuspended in 50 µL 8 M urea binding buffer (see above) and incubated at room temperature for 1 h to dissolve the protein pellet.

### Data presentation and statistical analysis

Boxplots were generated using GraphPad Prism 8.4.2 software (GraphPad software, Boston, MA, USA). Statistical analysis of quantitative data is based on ordinary one-way ANOVA followed by Tukey’s multiple comparison test (conducted in GraphPad Prism).

## Abbreviations

BiFC: Bimolecular fluorescence complementation
CAM: Calmodulin
CAMBD: Calmodulin binding domain
CML: Calmodulin-like
CT: Cytoplasmic C-terminus
EDTA: ethylenediamine-*N*,*N*,*N*′,*N*′-tetraacetic acid
EGTA: Ethylene glycol-bis(β-aminoethyl ether)-*N*,*N*,*N*′,*N*′-tetraacetic acid GST Glutathione *S*-transferase
HRP: Horseradish peroxidase
LCI: Luciferase complementation imaging
LUC: Luciferase
MLO: Mildew resistance locus o
Ni-NTA: Nickel nitrilotriacetic acid
OD: Optical density
PBS: Phosphate-buffered saline
Rpm: Revolutions per minute
SC: Synthetic complete
SDS: Sodium dodecyl sulfate
SDS-PAGE: Sodium dodecyl sulfate polyacrylamide gel electrophoresis SUS Split-ubiquitin system
TbID: TurboID biotin ligase
TBST: Tris-buffered saline with Tween20
Y2H: Yeast two-hybrid
WT: Wild-type

## Acknowledgments

Molecular graphics and analyses of the MLO2^CT^ three-dimensional protein structure were performed with UCSF ChimeraX, developed by the Resource for Biocomputing, Visualization, and Informatics at the University of California, San Francisco, with support from National Institutes of Health R01-GM129325 and the Office of Cyber Infrastructure and Computational Biology, National Institute of Allergy and Infectious Diseases. We thank Gitta Coaker (UC Davis, USA) for sharing pCAMBIA-based LCI vectors and Prof. Dr. Karin Römisch (Saarland University, Germany) for providing an aliquot of the α-LexA antiserum.

## Funding

This work was supported by grants of the Deutsche Forschungsgemeinschaft (DFG; PA861/20-1, project number 411779037) and the Novo Nordisk Foundation (grant NNF19OC0056457, PlantsGoImmune) to R.P. as well as the project “Grant Schemes at CU“ (reg. no. CZ.02.2.69/0.0/0.0/19_073/0016935) and a fund from the Czech Science Foundation/GACR 19-02242J awarded to A.B.F.

## Author contributions

K.B. generated constructs, performed the CAM overlay assays, the GST pull-down experiments, the BiFC assays, the LIC assays with MLO2^CT^ and the initial TbID assay with HA-CAM2.

B.S. generated constructs, performed the Y2H and yeast SUS assays as well as the TbID assay with LUC^C^-CAM2.

A.B.F. created the MLO2-TbID construct.

H.K. generated constructs and performed the LCI assays with full-length MLO2.

F.L. generated constructs, contributed to the conception of the study and supervised the practical work.

R.P. conceived the study, performed the *in silico* analyses and wrote the manuscript.

All authors read and commented on the manuscript.

## Supplemental files

Supplemental File 1. Relevant amino acid sequences.

**`>sp|Q9SXB6|MLO2_ARATH MLO-like protein 2 OS=Arabidopsis thaliana OX=3702 GN=MLO2 PE=1 SV=1 (C-terminus highlighted in yellow)`**

MADQVKERTLEETSTWAVAVVCFVLLFISIVLEHSIHKIGTWFKKKHKQALFEALEKVKA ELMLLGFISLLLTIGQTPISNICISQKVASTMHPCSAAEEAKKYGKKDAGKKDDGDGDKP GRRLLLELAESYIHRRSLATKGYDKCAEKGKVAFVSAYGIHQLHIFIFVLAVVHVVYCIV TYAFGKIKMRTWKSWEEETKTIEYQYSNDPERFRFARDTSFGRRHLNFWSKTRVTLWIVC FFRQFFGSVTKVDYLALRHGFIMAHFAPGNESRFDFRKYIQRSLEKDFKTVVEISPVIWF VAVLFLLTNSYGLRSYLWLPFIPLVVILIVGTKLEVIITKLGLRIQEKGDVVRGAPVVQP GDDLFWFGKPRFILFLIHLVLFTNAFQLAFFAWSTYEFNLNNCFHESTADVVIRLVVGAV VQILCSYVTLPLYALVTQMGSKMKPTVFNDRVATALKKWHHTAKNETKHGRHSGSNTPFS SRPTTPTHGSSPIHLLHNFNNRSVENYPSSPSPRYSGHGHHEHQFWDPESQHQEAETSTH HSLAHESSEPVLASVELPPIRTSKSLRDFSFKK

***`A. thaliana`* `MLO2 C-terminus (CAMBD highlighted in green)`** MGSKMKPTVFNDRVATALKKWHHTAKNETKHGRHSGSNTPFSSRPTTPTHGSSPIHLLHNF NNRSVENYPSSPSPRYSGHGHHEHQFWDPESQHQEAETSTHHSLAHESSEPVLASVELPPI RTSKSLRDFSFKK

**`>sp|P93766|MLO_HORVU Protein MLO OS=Hordeum vulgare OX=4513 GN=MLO PE=1 SV=1 (C-terminus highlighted in light blue)`**

MSDKKGVPARELPETPSWAVAVVFAAMVLVSVLMEHGLHKLGHWFQHRHKKALWEALEKM KAELMLVGFISLLLIVTQDPIIAKICISEDAADVMWPCKRGTEGRKPSKYVDYCPEGKVA LMSTGSLHQLHVFIFVLAVFHVTYSVITIALSRLKMRTWKKWETETTSLEYQFANDPARF RFTHQTSFVKRHLGLSSTPGIRWVVAFFRQFFRSVTKVDYLTLRAGFINAHLSQNSKFDF HKYIKRSMEDDFKVVVGISLPLWGVAILTLFLDINGVGTLIWISFIPLVILLCVGTKLEM IIMEMALEIQDRASVIKGAPVVEPSNKFFWFHRPDWVLFFIHLTLFQNAFQMAHFVWTVA TPGLKKCYHTQIGLSIMKVVVGLALQFLCSYMTFPLYALVTQMGSNMKRSIFDEQTSKAL TNWRNTAKEKKKVRDTDMLMAQMIGDATPSRGSSPMPSRGSSPVHLLHKGMGRSDDPQSA PTSPRTQQEARDMYPVVVAHPVHRLNPNDRRRSASSSALEADIPSADFSFSQG

**`Barley Mlo C-terminus (CAMBD highlighted in magenta)`** MGSNMKRSIFDEQTSKALTNWRNTAKEKKKVRDTDMLMAQMIGDATPSRGSSPMPSRGSSP VHLLHKGMGRSDDPQSAPTSPRTQQEARDMYPVVVAHPVHRLNPNDRRRSASSSALEADIP SADFSFSQG

**Amino acid sequence alignment of the CAMBDs of *A. thaliana* MLO2 and barley Mlo. Conserved amino acids are shown in green.**

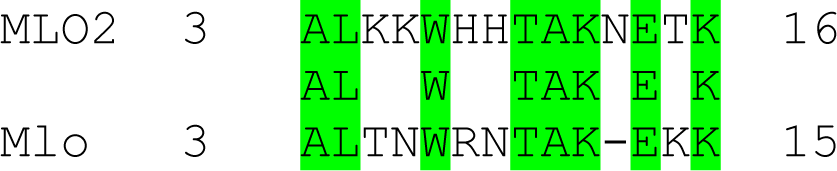

**Supplemental Figure 1.**
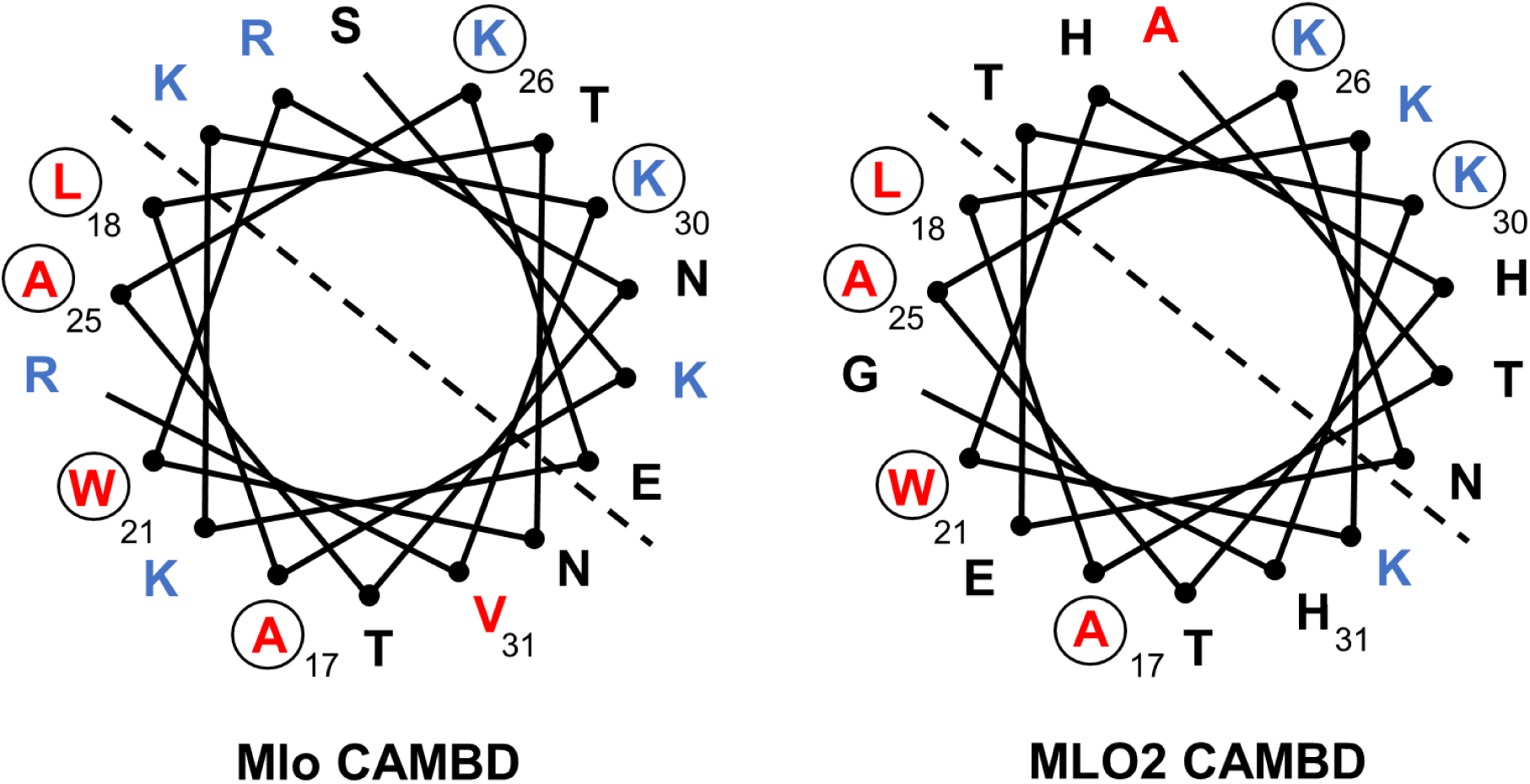
Conservation of amino acids in the barley Mlo and *A. thaliana* MLO2 CAMBDs. Helical wheel projections of the barley Mlo CAMBD (left; numbering according to the Mlo C-terminus; see **Supplemental File 1**) and *A. thaliana* MLO2 CAMBD (right; numbering according to the MLO2 C-terminus; see **Supplemental File 1**). Individual residues are indicated corresponding to the single-letter code for amino acids. The dashed line separates one side of the helix with preferentially hydrophobic residues (red; bottom left) from another side of the helix with preferentially basic residues (blue; top right). Relevant hydrophobic (red) and basic (blue) amino acid residues that reside in a conserved relative position between the barley Mlo CAMBD (left) and the *A. thaliana* MLO2 CAMBD (right) are marked with a circle.

**Supplemental Figure 2.**
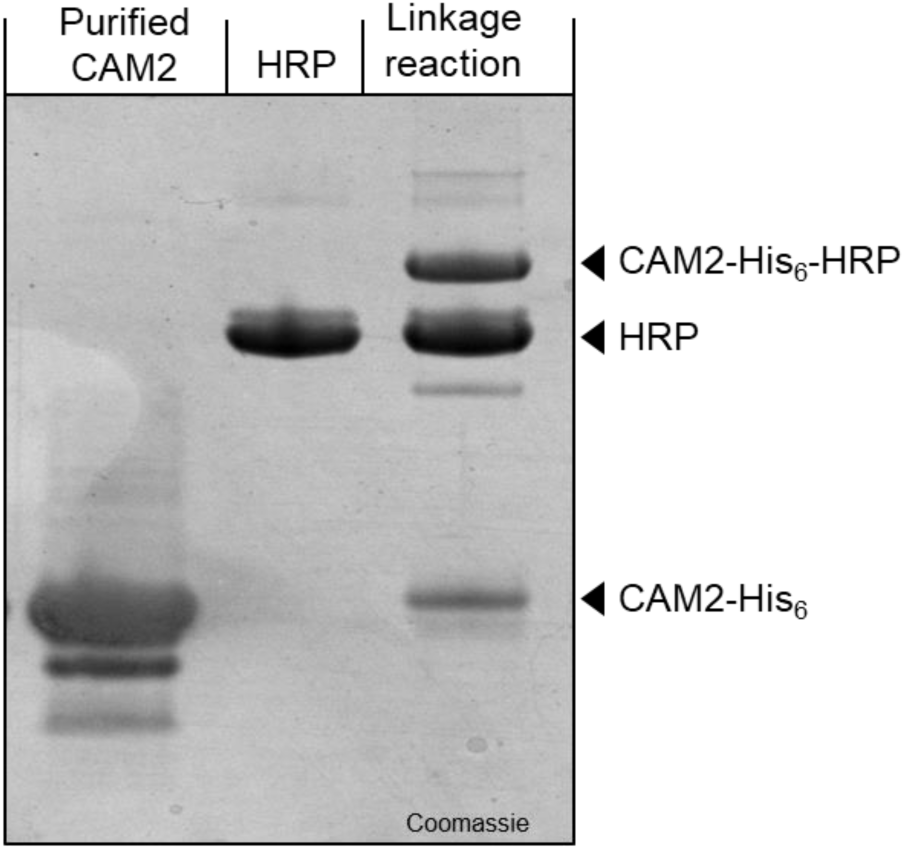
Chemical linkage of CAM2 to HRP. SDS-PAGE illustrating the efficiency of the chemical linkage reaction between affinity-purified CAM2 (reduced in 0.5 mM TCEP prior to gel loading) and maleimide-coupled HRP. Expected molecular masses of CAM2-His_6_, HRP and the CAM2-His_6_-HRP conjugation product are marked by a black triangle. The gel was stained with Coomassie Brilliant Blue solution.

**Supplemental Figure 3.**
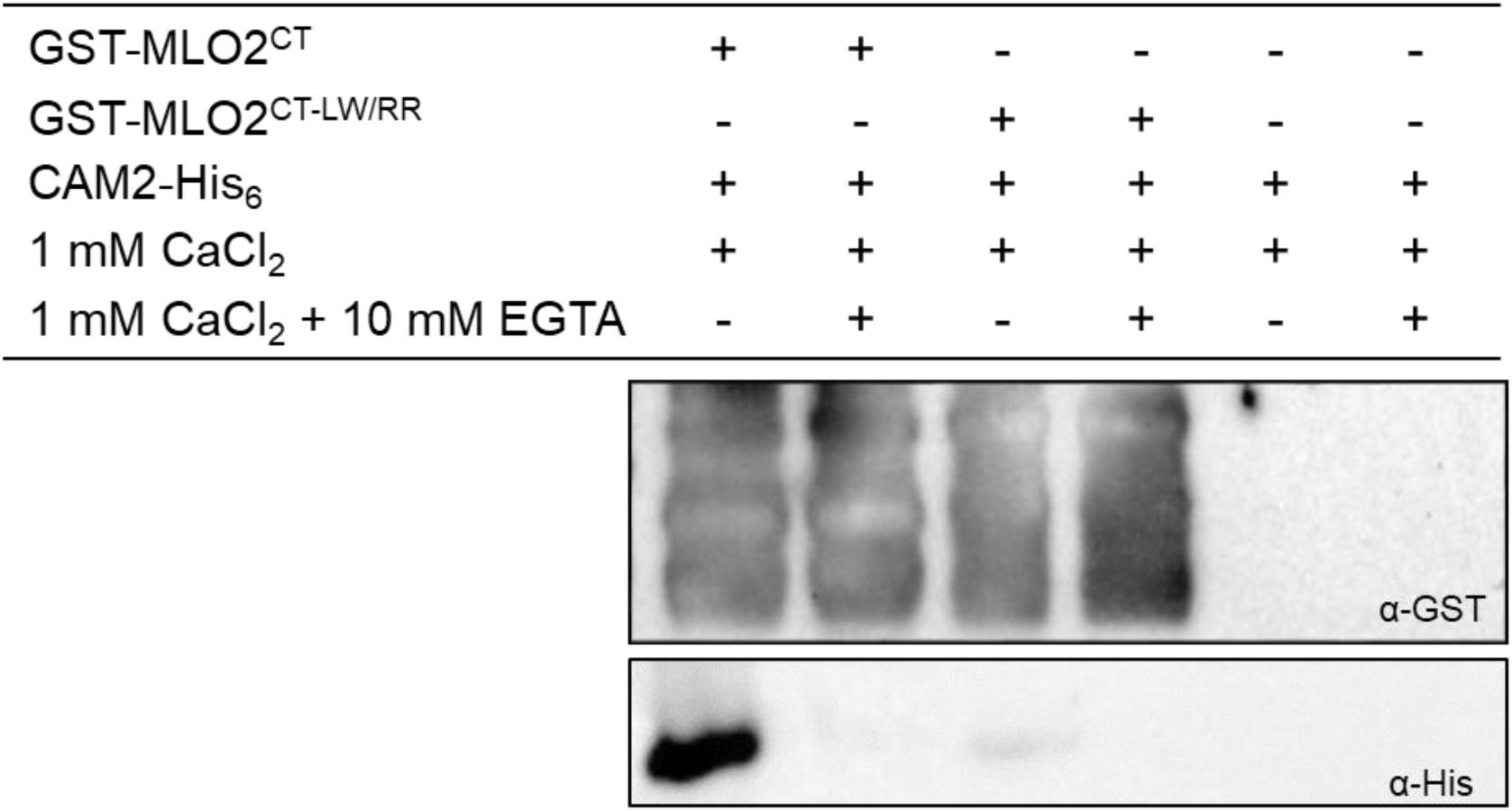
Initial GST pull-down assay. GST pulldown assay with recombinantly expressed GST-tagged MLO2^CT^ and MLO2^CT-LW/RR^ plus CAM-His_6_ in the presence of either 1 mM CaCl_2_ or 1 mM CaCl_2_ plus 10 mM EGTA. Protein input was assessed by immunoblot analysis with an α-GST antibody (upper panel). Presence of CAM-His_6_ was analyzed by immunoblot analysis with an α-His antibody (lower panel).

**Supplemental Figure 4.**
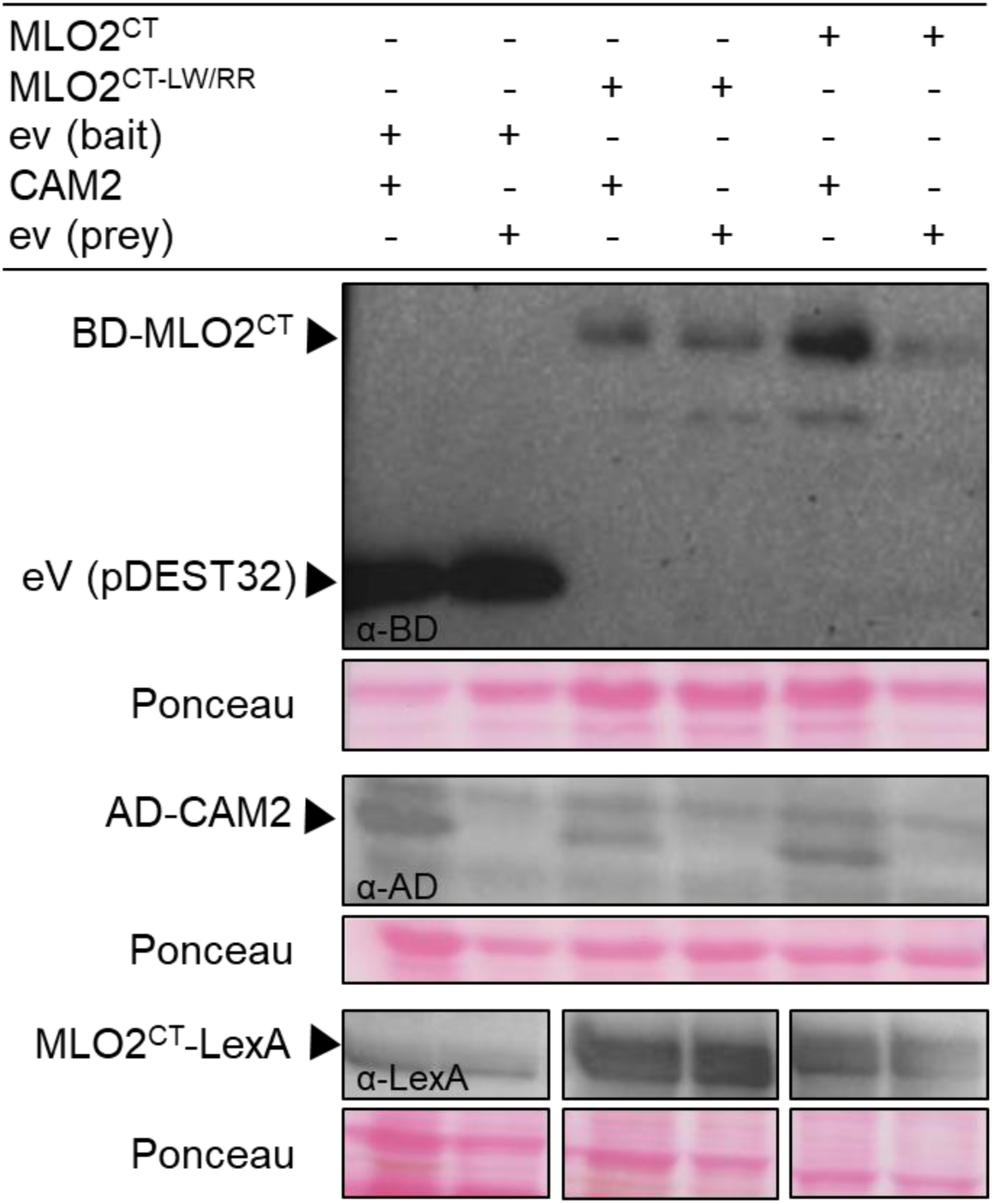
Immunoblot analysis for the pDEST32/pDEST22-based Y2H and PLV-based yeast SUS assays. Yeast protein extracts were prepared, separated by SDS-PAGE, blotted on nitrocellulose membrane and subjected to immunodetection using specific antibodies. For the pDEST32/pDEST22-based Y2H assay, blots were probed with α-BD and α-AD-specific antibodies for the detection of MLO2^CT^, MLO2^CT-LW/RR^ and CAM2 (upper panels). For the PLV-based yeast SUS based on the yeast Ost4 membrane protein, the blot was probed with an α-LexA-specific antibody for the detection of Ost4-fused MLO2^CT^ and MLO2^CT-LW/RR^ (lower panel). Empty vector (ev) controls were included for both types of assays. Ponceau staining served in all cases to judge equal gel loading.

**Supplemental Figure 5.**
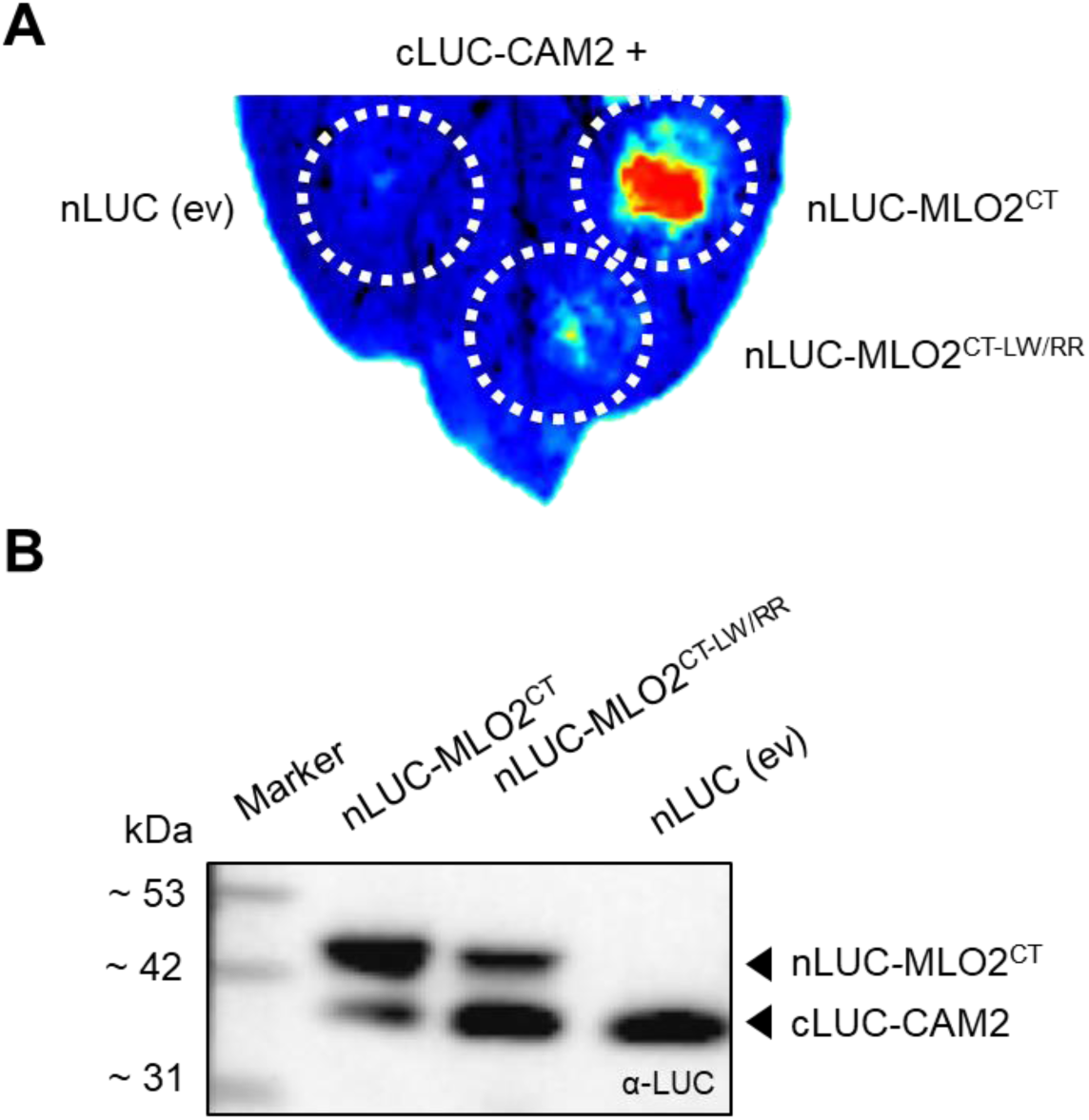
Representative leaf and immunoblot analysis related to the LCI assay. **A** Representative *N. benthamiana* leaf showing luciferase-based luminescence upon transient expression of the indicated construct combinations. Circled lines indicate the sites of agrobacteria infiltration. An empty vector (ev) control was included for nLUC. **B** Immunoblot analysis of *N. benthamiana* leaf extracts upon transient expression of the indicated construct combinations. The blot was probed with an α-LUC antibody.

**Supplemental Figure 6.**
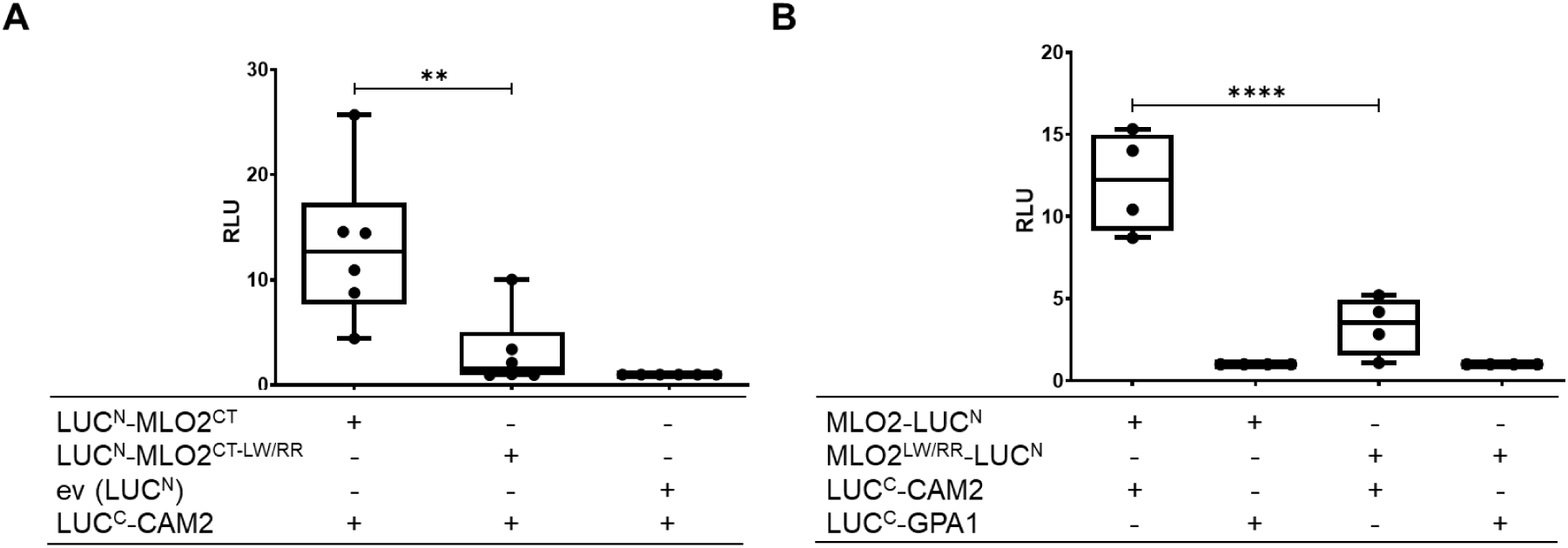
Representation of LCI assay data by relative light units. Data shown for the LCI assay in Figure 7 were normalized to the respective negative controls, i.e. empty vector (eV) in the case of MLO2^CT^ variants (**A**) and LUC^C^-GPA1 in case of the MLO full-length variants (**B**). Normalized values are given as relative light units (RLU). Asterisks indicate a statistically significant difference between MLO2^CT^ or MLO2 in comparison to the respective LW/RR double mutant variant according to one-way ANOVA followed by Tukey’s multiple comparison test (** *p*<0.01, **** *p*<0.0001).

**Supplemental Table 1.** Media composition for yeast-based interaction assays.

| Yeast system | Growth control medium | Interaction-selective medium |
| --- | --- | --- |
| classical Y2H (Invitrogen) | SC-Leu-Trp | SC-Leu-Trp-His |
| classical Y2H (Clontech) | SC-Leu-Trp | SC-Leu-Trp-His |
| Ura3-based yeast SUS | SC-His-Trp | SC-His-Trp + 0.7 g/L 5-FOA |
| PLV-based yeast SUS | SC-Leu-Trp | SC-Leu-Trp-Ade<br>SC-Leu-Trp + 10 mM 3-AT |

## Notes

### Competing Interest Statement

The authors have declared no competing interest.

## References

Arora, D., Abel, N.B., Liu, C., van Damme, P., Yperman, K., Eeckhout, D., et al. (2020) Establishment of proximity-dependent biotinylation approaches in different plant model systems. Plant Cell, 32: 3388–3407.

Bhat, R.A., Miklis, M., Schmelzer, E., Schulze-Lefert, P., and Panstruga, R. (2005) Recruitment and interaction dynamics of plant penetration resistance components in a plasma membrane microdomain. Proceedings of the National Academy of Sciences of the United States of America, 102: 3135–3140.

Bidzinski, P., Noir, S., Shahi, S., Reinstädler, A., Gratkowska, D.M., and Panstruga, R. (2014) Physiological characterization and genetic modifiers of aberrant root thigmomorphogenesis in mutants of *Arabidopsis thaliana MILDEW LOCUS O* genes. *Plant*, Cell & Environment, 37: 2738–2753.

Boeke, J.D., Trueheart, J., Natsoulis, G., and Fink, G.R. (1987) 5-Fluoroorotic acid as a selective agent in yeast molecular genetics. In Recombinant DNA. Wu, R. (ed). San Diego, Calif.: Academic Press, pp. 164–175.

Bradford, M.M. (1976) A rapid and sensitive method for the quantitation of microgram quantities of protein utilizing the principle of protein-dye binding. Analytical Biochemistry, 72: 248–254.

Branon, T.C., Bosch, J.A., Sanchez, A.D., Udeshi, N.D., Svinkina, T., Carr, S.A., et al. (2018) Efficient proximity labeling in living cells and organisms with TurboID. Nature Biotechnology, 36: 880–887.

Campe, R., Langenbach, C., Leissing, F., Popescu, G.V., Popescu, S.C., Goellner, K., Beckers, Gerold J M, and Conrath, U. (2016) ABC transporter PEN3/PDR8/ABCG36 interacts with calmodulin that, like PEN3, is required for Arabidopsis nonhost resistance. New Phytologist, 209: 294–306.

Chen, H.M., Zou, Y., Shang, Y.L., Lin, H.Q., Wang, Y.J., Cai, R., Tang, X.Y., and Zhou, J.M. (2008) Firefly luciferase complementation imaging assay for protein-protein interactions in plants. Plant Physiology, 146: 368–376.

Chen, Z.Y., Noir, S., Kwaaitaal, M., Hartmann, H.A., Wu, M.J., Mudgil, Y., et al. (2009) Two seven-transmembrane domain MILDEW RESISTANCE LOCUS O proteins cofunction in *Arabidopsis* root thigmomorphogenesis. Plant Cell, 21: 1972–1991.

Consonni, C., Bednarek, P., Humphry, M., Francocci, F., Ferrari, S., Harzen, A., van Themaat, E.V.L., and Panstruga, R. (2010) Tryptophan-derived metabolites are required for antifungal defense in the Arabidopsis *mlo2* mutant. Plant Physiology, 152: 1544–1561.

Consonni, C., Humphry, M.E., Hartmann, H.A., Livaja, M., Durner, J., Westphal, L., et al. (2006) Conserved requirement for a plant host cell protein in powdery mildew pathogenesis. Nature Genetics, 38: 716–720.

Cox, J.S., Chapman, R.E., and Walter, P. (1997) The unfolded protein response coordinates the production of endoplasmic reticulum protein and endoplasmic reticulum membrane. Molecular Biology of the Cell, 8: 1805–1814.

Cui, F., Wu, H., Safronov, O., Zhang, P., Kumar, R., Kollist, H., et al. (2018) Arabidopsis MLO2 is a negative regulator of sensitivity to extracellular reactive oxygen species. Plant, Cell & Environment, 41: 782–796.

Deslandes, L., Olivier, J., Peeters, N., Feng, D.X., Khounlotham, M., Boucher, C., et al. (2003) Physical interaction between RRS1-R, a protein conferring resistance to bacterial wilt, and PopP2, a type III effector targeted to the plant nucleus. Proceedings of the National Academy of Sciences of the United States of America, 100: 8024–8029.

Devoto, A., Hartmann, H.A., Piffanelli, P., Elliott, C., Simmons, C., Taramino, G., et al. (2003) Molecular phylogeny and evolution of the plant-specific seven-transmembrane MLO family. Journal of Molecular Evolution, 56: 77–88.

Devoto, A., Piffanelli, P., Nilsson, I., Wallin, E., Panstruga, R., Heijne, G. von, and Schulze-Lefert, P. (1999) Topology, subcellular localization, and sequence diversity of the Mlo family in plants. Journal of Biological Chemistry, 274: 34993– 35004.

Dohmen, R.J., Stappen, R., McGrath, J.P., Forrová, H., Kolarov, J., Goffeau, A., and Varshavsky, A. (1995) An essential yeast gene encoding a homolog of ubiquitin-activating enzyme. Journal of Biological Chemistry, 270: 18099–18109.

Duan, G., and Walther, D. (2015) The roles of post-translational modifications in the context of protein interaction networks. PLoS Computational Biology, 11: e1004049.

Earley, K.W., Haag, J.R., Pontes, O., Opper, K., Juehne, T., Song, K., and Pikaard, C.S. (2006) Gateway-compatible vectors for plant functional genomics and proteomics. Plant Journal, 45: 616–629.

Ebel, C. (2007) Solvent mediated protein–protein interactions. In Protein interactions. Biophysical approaches for the study of complex reversible systems. Schuck, P. (ed). New York, NY: Springer, pp. 255–287.

Elliott, C., Müller, J., Miklis, M., Bhat, R.A., Schulze-Lefert, P., and Panstruga, R. (2005) Conserved extracellular cysteine residues and cytoplasmic loop-loop interplay are required for functionality of the heptahelical MLO protein. Biochemical Journal, 385: 243–254.

Erijman, A., Rosenthal, E., and Shifman, J.M. (2014) How structure defines affinity in protein-protein interactions. PLoS One, 9: e110085.

Gao, Q., Wang, C., Xi, Y., Shao, Q., Li, L., and Luan, S. (2022) A receptor–channel trio conducts Ca^2+^ signalling for pollen tube reception. Nature, 607: 534–539.

Gibson, D.G., Young, L., Chuang, R.-Y., Venter, J.C., Hutchison, C.A., and Smith, H.O. (2009) Enzymatic assembly of DNA molecules up to several hundred kilobases. Nature Methods, 6: 343–345.

Gietz, R.D., and Woods, R.A. (2002) Transformation of yeast by lithium acetate/single-stranded carrier DNA/polyethylene glycol method. In Guide to yeast genetics and molecular and cell biology. Fink, G.R., and Guthrie, C. (eds). Amsterdam: Academic Pr, pp. 87–96.

Grefen, C., Donald, N., Hashimoto, K., Kudla, J., Schumacher, K., and Blatt, M.R. (2010) A ubiquitin-10 promoter-based vector set for fluorescent protein tagging facilitates temporal stability and native protein distribution in transient and stable expression studies. Plant Journal, 64: 355–365.

Grefen, C., Obrdlik, P., and Harter, K. (2009) The determination of protein-protein interactions by the mating-based split-ubiquitin system (mbSUS). Methods in Molecular Biology, 479: 217–233.

Gruner, K., Leissing, F., Sinitski, D., Thieron, H., Axstmann, C., Baumgarten, K., et al. (2021) Chemokine-like MDL proteins modulate flowering time and innate immunity in plants. Journal of Biological Chemistry, 296: 100611.

Harty, C., and Römisch, K. (2013) Analysis of Sec61p and Ssh1p interactions in the ER membrane using the split-ubiquitin system. BMC Cell Biology, 14: 14.

Huebbers, J.W., Caldarescu, G.A., Kubátová, Z., Sabol, P., Levecque, S.C.J., Kuhn, H., et al. (2022) Interplay of EXO70 and MLO proteins modulates trichome cell wall composition and powdery mildew susceptibility. bioRxiv.

James, P., Halladay, J., and Craig, E.A. (1996) Genomic libraries and a host strain designed for highly efficient two-hybrid selection in yeast. Genetics, 144: 1425– 1436.

Johnsson, N., and Varshavsky, A. (1994) Split ubiquitin as a sensor of protein interactions in vivo. Proceedings of the National Academy of Sciences of the United States of America, 91: 10340–10344.

Jones, D.S., Yuan, J., Smith, B.E., Willoughby, A.C., Kumimoto, E.L., and Kessler, S.A. (2017) MILDEW RESISTANCE LOCUS O function in pollen tube reception is linked to its oligomerization and subcellular sistribution. Plant Physiology, 175: 172–185.

Jørgensen, J.H. (1992) Discovery, characterization and exploitation of Mlo powdery mildew resistance in barley. Euphytica, 63: 141–152.

Jumper, J., Evans, R., Pritzel, A., Green, T., Figurnov, M., Ronneberger, O., et al. (2021) Highly accurate protein structure prediction with AlphaFold. Nature, 596: 583–589.

Karimi, M., Meyer, B. de, and Hilson, P. (2005) Modular cloning in plant cells. Trends in Plant Science, 10: 103–105.

Kessler, S.A., Shimosato-Asano, H., Keinath, N.F., Wuest, S.E., Ingram, G., Panstruga, R., and Grossniklaus, U. (2010) Conserved molecular components for pollen tube reception and fungal invasion. Science, 330: 968–971.

Kim, D.S., Choi, H.W., and Hwang, B.K. (2014) Pepper mildew resistance locus O interacts with pepper calmodulin and suppresses Xanthomonas AvrBsT-triggered cell death and defense responses. Planta, 240: 827–839.

Kim, M.C., Lee, S.H., Kim, J.K., Chun, H.J., Choi, M.S., Chung, W.S., et al. (2002a) Mlo, a modulator of plant defense and cell death, is a novel calmodulin-binding protein - Isolation and characterization of a rice Mlo homologue. Journal of Biological Chemistry, 277: 19304–19314.

Kim, M.C., Panstruga, R., Elliott, C., Müller, J., Devoto, A., Yoon, H.W., et al. (2002b) Calmodulin interacts with MLO protein to regulate defence against mildew in barley. Nature, 416: 447–450.

Kudla, J., and Bock, R. (2016) Lighting the way to protein-protein interactions: Recommendations on best practices for Bimolecular Fluorescence Complementation analyses. Plant Cell, 28: 1002–1008.

Kusch, S., and Panstruga, R. (2017) *mlo*-based resistance: An apparently universal “weapon” to defeat powdery mildew disease. Molecular Plant-Microbe Interactions, 30: 179–189.

Kusch, S., Pesch, L., and Panstruga, R. (2016) Comprehensive phylogenetic analysis sheds light on the diversity and origin of the MLO family of integral membrane proteins. Genome Biology and Evolution, 8: 878–895.

Kusch, S., Thiery, S., Reinstädler, A., Gruner, K., Zienkiewicz, K., Feussner, I., and Panstruga, R. (2019) Arabidopsis *mlo3* mutant plants exhibit spontaneous callose deposition and signs of early leaf senescence. Plant Molecular Biology, 101: 21– 40.

Lyngkjær, M.F., Newton, A.C., Atzema, J.L., and Baker, S.J. (2000) The barley *mlo* gene: an important powdery mildew resistance source. Agronomie, 20: 745–756.

Mair, A., Xu, S.-L., Branon, T.C., Ting, A.Y., and Bergmann, D.C. (2019) Proximity labeling of protein complexes and cell-type-specific organellar proteomes in *Arabidopsis* enabled by TurboID. eLife, 8.

McCormack, E., Tsai, Y.-C., and Braam, J. (2005) Handling calcium signaling: Arabidopsis CaMs and CMLs. Trends in Plant Science, 10: 383–389.

Meng, J.-G., Liang, L., Jia, P.-F., Wang, Y.-C., Li, H.-J., and Yang, W.-C. (2020) Integration of ovular signals and exocytosis of a Ca^2+^ channel by MLOs in pollen tube guidance. Nature Plants, 6: 143–153.

Miller, K.E., Kim, Y., Huh, W.-K., and Park, H.-O. (2015) Bimolecular Fluorescence Complementation (BiFC) analysis: Advances and recent applications for genome-wide interaction studies. Journal of Molecular Biology, 427: 2039–2055.

Möckli, N., Deplazes, A., Hassa, P.O., Zhang, Z., Peter, M., Hottiger, M.O., Stagljar, I., and Auerbach, D. (2007) Yeast split-ubiquitin-based cytosolic screening system to detect interactions between transcriptionally active proteins. BioTechniques, 42: 725–730.

Obrdlik, P., El-Bakkoury, M., Hamacher, T., Cappellaro, C., Vilarino, C., Fleischer, C., et al. (2004) K^+^ channel interactions detected by a genetic system optimized for systematic studies of membrane protein interactions. Proceedings of the National Academy of Sciences of the United States of America, 101: 12242–12247.

Panstruga, R. (2005) Discovery of novel conserved peptide domains by ortholog comparison within plant multi-protein families. Plant Molecular Biology, 59: 485– 500.

Pettersen, E.F., Goddard, T.D., Huang, C.C., Meng, E.C., Couch, G.S., Croll, T.I., Morris, J.H., and Ferrin, T.E. (2021) UCSF ChimeraX: Structure visualization for researchers, educators, and developers. Protein Science, 30: 70–82.

Piffanelli, P., Devoto, A., and Schulze-Lefert, P. (1999) Defence signalling pathways in cereals. Current Opinion in Plant Biology, 2: 295–300.

Samalova, M., Brzobohaty, B., and Moore, I. (2005) pOp6/LhGR: A stringently regulated and highly responsive dexamethasone-inducible gene expression system for tobacco. Plant Journal, 41: 919–935.

Schütze, K., Harter, K., and Chaban, C. (2009) Bimolecular fluorescence complementation (BiFC) to study protein-protein interactions in living plant cells. Methods in Molecular Biology, 479: 189–202.

Stagljar, I., Korostensky, C., Johnsson, N., and te Heesen, S. (1998) A genetic system based on split-ubiquitin for the analysis of interactions between membrane proteins in vivo. Proceedings of the National Academy of Sciences of the United States of America, 95: 5187–5192.

Walter, M., Chaban, C., Schutze, K., Batistic, O., Weckermann, K., Nake, C., et al. (2004) Visualization of protein interactions in living plant cells using bimolecular fluorescence complementation. Plant Journal, 40: 428–438.

Wittke, S., Lewke, N., Müller, S., and Johnsson, N. (1999) Probing the molecular environment of membrane proteins in vivo. Molecular Biology of the Cell, 10: 2519–2530.

Xing, S., Wallmeroth, N., Berendzen, K.W., and Grefen, C. (2016) Techniques for the analysis of protein-protein interactions in vivo. Plant Physiology, 171: 727–758.

Xue, B., Dunbrack, R.L., Williams, R.W., Dunker, A.K., and Uversky, V.N. (2010) PONDR-FIT: A meta-predictor of intrinsically disordered amino acids. Biochimica Et Biophysica Acta, 1804: 996–1010.

Yang, X., Wen, Z., Zhang, D., Li, Z., Li, D., Nagalakshmi, U., Dinesh-Kumar, S.P., and Zhang, Y. (2021) Proximity labeling: an emerging tool for probing *in planta* molecular interactions. Plant Communications, 2: 100137.

Yu, G., Wang, X., Chen, Q., Cui, N., Yu, Y., and Fan, H. (2019) Cucumber Mildew Resistance Locus O interacts with calmodulin and regulates plant cell death associated with plant immunity. International Journal of Molecular Sciences, 20.

Zhang, Y., Li, Y., Yang, X., Wen, Z., Nagalakshmi, U., and Dinesh-Kumar, S.P. (2020) TurboID-based proximity labeling for in planta identification of protein-protein interaction networks. Journal of Visualized Experiments : JoVE.

Zhu, L., Zhang, X.-Q., de Ye, and Chen, L.-Q. (2021) The Mildew Resistance Locus O 4 interacts with CaM/CML and is involved in root gravity response. International Journal of Molecular Sciences, 22: 5962.

